# MAPseq2: a sensitive barcoded connectomics method

**DOI:** 10.1101/2025.06.23.661165

**Authors:** Hyopil Kim, Huihui Qi, Craig Washington, Xiaoping Liang, Justus M. Kebschull

## Abstract

The barcoded connectomics tool MAPseq enables highly multiplexed projection mapping of individual neurons by translating neuroanatomy into a format solvable by DNA sequencing. Here we present MAPseq2, a user-friendly protocol with up to ∼10-fold increased sensitivity compared to the current state of the art and ∼10-fold decreased cost. As MAPseq workflows are used across a range of barcoded connectomics methods, including BARseq, BRICseq, and ConnectID, all improvements in MAPseq2 directly transfer to these technologies.

## Main

Individual neurons exhibit heterogeneous projections to distant brain regions.^1,2^ Determining the structure of these projections is critical for deciphering information flow in the vertebrate brain.^3,4^ High-resolution imaging-based methods including fMOST,^5,6^ serial 2-photon tomography,^7,8^ and expansion-assisted light sheet imaging^9–11^ have been used to map brain-wide single-cell projections in mammalian brains. However, these methods remain technically demanding and labor-intensive. Moreover, they struggle to connect projections to the transcriptomic identity of individual neurons.

To overcome these limitations, we previously developed the first-in-class barcoded connectomics method MAPseq^12^ (Fig. 1a). In MAPseq, thousands of neurons in a source region are uniquely labelled with barcodes delivered by a viral library. The barcodes are expressed as mRNAs and transported to axon terminals by a carrier protein. Barcode mRNAs from the injection site and potential axonal target sites are then sequenced and matched across sites to reconstruct thousands of single-neuron projection patterns in single animals.

**Figure 1.**
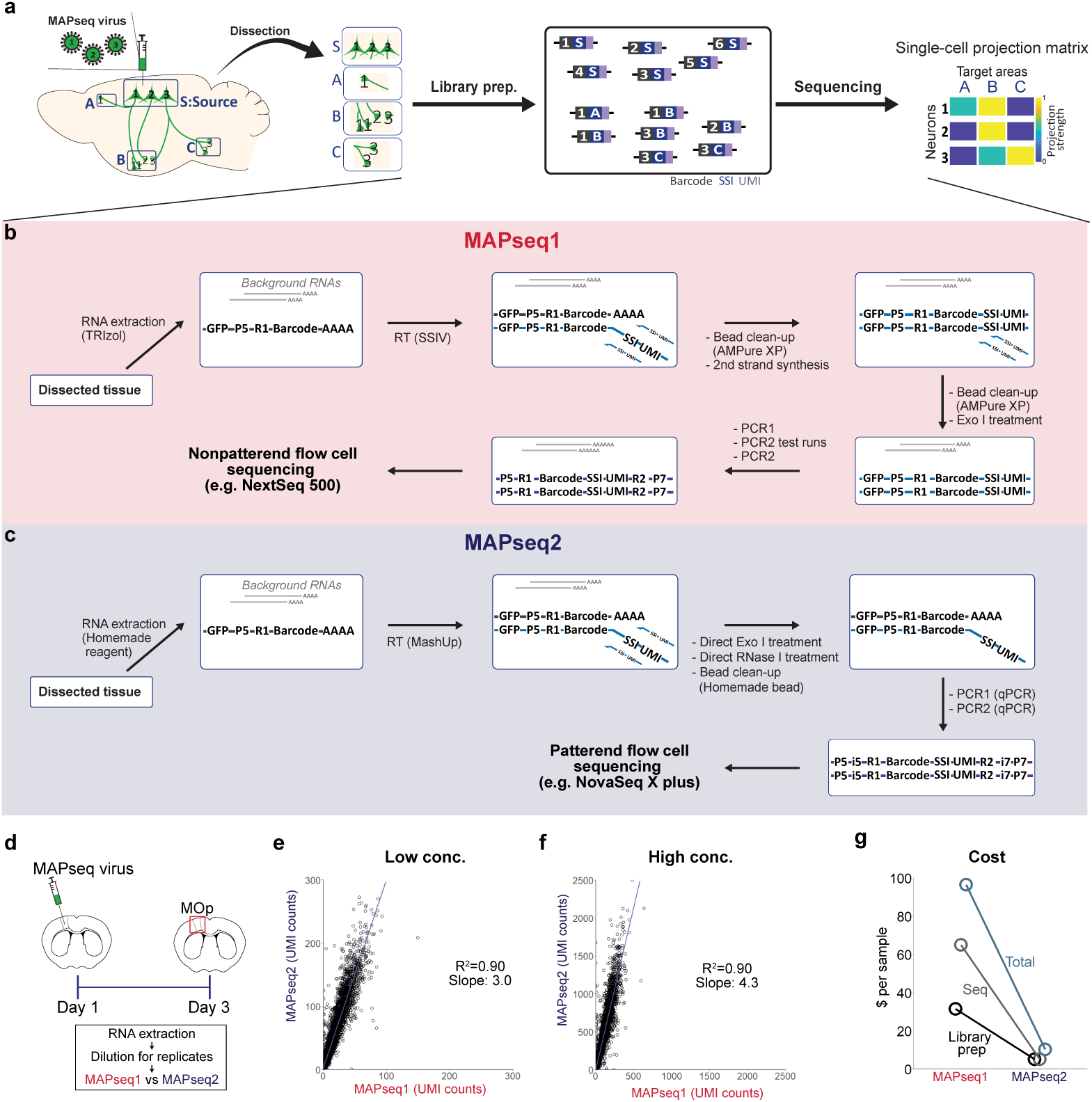
MAPseq2 increases sensitivity, decreases costs, and simplifies the MAPseq workflow. **a,** Schematic of the MAPseq workflow. Barcoded RNAs extracted from target regions are converted into sequencing libraries and sequenced to reconstruct single-neuron projection matrices. Numbers and letters represent unique barcodes and sample-specific indices (SSIs), respectively. **b, c,** Library preparation and sequencing workflows of MAPseq1 (**b**) and MAPseq2 (**c**). **d-f,** Evaluation of MAPseq2 barcode recovery compared to MAPseq1. **d,** Experimental design where MAPseq virus was injected into mouse primary motor cortex and RNA was extracted from the injection site. 100 times diluted low barcode concentration samples (**e**) and 10 times diluted high barcode concentration samples (**f**) were subjected to MAPseq1 and MAPseq2 (N=3). **e, f,** Comparison of barcode recovery between MAPseq1 and MAPseq2 using the low (**e**) and high (**f**) concentration samples. MAPseq2 recovered more barcodes than MAPseq1 with high correlation scores. Each dot represents a barcode and mean UMI values from three replicates are plotted (**e,** linear regression, R^2^=0.90, slope=3.0, *p*<0.0001; **f,** linear regression, R^2^=0.90, slope=4.3, *p*<0.0001). **g,** Comparison of cost per sample assuming 10 million reads. MAPseq2 reduces library prep cost by 6.2-fold ($31.50 to $5.10), sequencing cost by 12.6-fold ($65.00 to $5.20), and total cost by 9.4-fold ($96.50 to $10.30).

A variety of methods now build on MAPseq, including ConnectID,^13^ BARseq,^14,15^, BRICseq,^16^ and POINTseq.^17^ In ConnectID and BARseq, barcoded cell bodies are subjected to single-cell RNA sequencing and in situ sequencing, respectively, to link projections to gene expression of thousands of individual cells. In BRICseq, hundreds of source sites are mapped simultaneously to enable a network level mapping of hundreds of thousands of cells and POINTseq enables cell type specific barcoding. MAPseq and its derivatives have been used and validated in various brain regions and species, including locus coeruleus, cerebral cortex, olfactory bulb, and hippocampus in mice, cortex in costarican singing mice, and the amygdala in macaque.^4,12,14–16,18–22^ While these methods have expanded the utility of barcoded connectomics, importantly, they all use the same library preparation and sequencing workflows as MAPseq whether for target sites in ConnectID and BARseq, or for the hundreds of samples in BRICseq.

Despite the high throughput of MAPseq, the original protocol (MAPseq1) remains inefficient, complicated, lengthy, and expensive, hindering the widespread use of barcoded connectomics (Fig. 1b; Supplementary Fig. 1a). MAPseq1 detects only 1–5% of barcode mRNAs and struggles with high RNA input, decreasing its ability to detect weak projections and compressing the dynamic range of the measurement.^12^ The protocol is reagent intensive, involving second strand cDNA synthesis and several clean-up reactions,^12^ and its sequencing amplicon structure is susceptible to index hopping on patterned Illumina flow cells,^23^ restricting MAPseq1 sequencing to older generation sequencers. These factors combine to drive up the cost of barcoded connectomics and limit the scale at which these experiments can be conducted. To address these limitations, we here developed MAPseq2 as a more sensitive method with lower reagent and sequencing costs and a streamlined workflow.

In MAPseq1 (Fig. 1b; Supplementary Fig. 1a), total RNA is extracted from all brain regions of interest. RNA barcodes are then reverse transcribed using SuperScript IV reverse transcriptase and gene specific primers, which include a sample-specific index (SSI) and a unique molecular identifier (UMI). As residual reverse transcription (RT) primers impede barcode quantification by creating new barcode-UMI associations during subsequent PCR steps, barcode cDNA is double-stranded by second strand cDNA synthesis, and single-stranded residual RT primers are removed by single-strand DNA specific Exonuclease I (Exo I) digestion. These steps involve two rounds of Ampure XP bead clean-up. The barcode fragments are then amplified by two rounds of PCR, where optimal cycle numbers are empirically determined by test runs of endpoint PCR to minimize PCR errors and template switches^24,25^. The amplicons are then sequenced on older generation Illumina machines with traditional flow cells to avoid index-hopping events on recently developed patterned flow cells^23^ that can create false-positive projections by switching barcode-SSI associations.

To reduce costs and hands-on time of MAPseq1, multiple samples are sometimes pooled after RT to be processed together in one reaction for subsequent steps.^16^ While this strategy reduces hands on work and provides some cost savings, it risks PCR template switching between samples, which produces artifactual barcode-SSI associations and thus false positive projections.^24^ Such PCR template switching can be reduced by increasing PCR volumes, as done in BRICseq, but this loses many of the practical advantages of sample pooling.^16^ Template switched reads can also be partially removed computationally.^16^ However, any sample pooling strategy fundamentally carries a significant risk of generating false-positive projections.

In MAPseq2 (Fig. 1c; Supplementary Fig. 1b), we did not use any sample pooling to completely avoid the risk of false positives from PCR template switching and optimized each of the steps in the MAPseq1 protocol to increase efficiency. We first redesigned the MAPseq sequencing amplicon structure to include matched pairs of i5 and i7 sample indexes. This change enables us to identify and discard index hopping events as reads with mismatched indexes and thus allows us to safely use the latest sequencing technologies^23^. Importantly, we chose to maintain the SSI sequences in RT primers to detect and eliminate sample cross-contamination after RT but before PCR2. We also optimized RT and PCR1 primers to minimize non-specific amplification (Supplementary Fig. 2a). We then streamlined the experimental protocol. Here we sought to eliminate second-strand synthesis and subsequent clean-up steps to limit barcode loss. We reasoned that the RNA-cDNA hybrid formed after 1^st^ strand cDNA synthesis ought to be sufficient to protect barcode cDNA from ExoI digestion, and that the unpaired RT primer overhangs should to be unaffected as Exo I only acts in the 3’ to 5’ direction.^26^ Indeed, Exo I efficiently removes RT primers without digesting the first strand cDNA when added to the RT reaction after extension but before denaturation (Supplementary Fig. 2b, c). We also found that background RNA carried over from RT interferes with PCR (Supplementary Fig. 2d). We therefore introduced an RNase I treatment after ExoI digestion to remove RNA before PCR and effectively rescued PCR efficiency with comparable Cq values across a wide range of background RNA (Supplementary Fig. 2e-g). Finally, we eliminated the need for test endpoint PCRs to determine the optimal number of PCR cycle numbers by monitoring amplification in real-time in the final PCR reaction using SYBR green quantitative PCR, improving our ability to minimize PCR induced deviations in our sequencing libraries^24^ and dramatically reducing reagent usage and hands-on time.^25^

In addition to these protocol design changes, we optimized reagents that are usually off the shelf. We tested efficiency improvements from different formulations of TRIzol reagent, RT enzymes, and cleanup beads. Importantly, we found that while the standard formulation for a home-made TRIzol reagent^27,28^ performs equivalently to the commercial reagent, further optimization of the homemade reagent yielded 3.2-fold higher RNA levels than the commercial TRIzol used in MAPseq1 from fresh-frozen brain tissues (Supplementary Fig. 2h). We also found that the open-source RT enzyme MashUp^29,30^ can substitute for SuperscriptIV (Supplementary Fig. 2i), and homemade cleanup beads prepared from Sera-Mag Magnetic SpeedBeads^31^ can substitute from Ampure XP beads (Supplementary Fig. 2j). Although the efficiency of MashUp and homemade beads was comparable to the commercial products, these changes yielded significant cost savings (Supplementary Table 1).

To directly compare the efficiency of MAPseq2 to MAPseq1 in recovering barcodes from RNA, we injected barcoded MAPseq virus^32^ into the mouse primary motor cortex and extracted total RNA from the injection site using commercial TRIzol reagent. We then diluted and split this RNA into replicate samples with high and low barcode concentration (N=3 each), prepared libraries of each sample using the two protocols, and sequenced libraries to full depth^12^ (Fig. 1d, Supplementary Fig. 3). Barcode abundances measured by MAPseq1 and MAPseq2 were highly correlated across (Fig. 1e,f) and within protocols (Supplementary Fig. 4), demonstrating the reliability and validity of our new protocol. Importantly, MAPseq2 recovered barcodes at 3 and 4.3 fold higher rates than MAPseq1 in the low and high concentration samples, respectively, demonstrating an increased sensitivity of MAPseq2 over MAPseq1 when starting with identical RNA input. Taken together with the 3.2-fold enhanced RNA extraction of the optimized home- made TRIzol, MAPseq2 can achieve up to ∼10 times increased barcode detection sensitivity over the previous MAPseq1 state of the art. Moreover, the differential performance of the two methods in high vs low concentration samples also indicates that MAPseq2 has a higher capacity for barcode detection per sample (Supplementary Fig. 5 and associated Supplementary Note). In addition to these increases in efficiency, MAPseq2 reduces library preparation costs by up to 6.2 times and sequencing costs by up 12.6 times. Assuming ∼10M reads per sample, this reduces overall MAPseq cost by ∼10-fold to $10.30 per sample (Fig. 1g; Supplementary Table 1).

To validate MAPseq2 in mapping brain-wide projections patterns, we focused on mouse primary motor cortex. Primary motor cortex has been extensively studied,^20,33,34^ including using MAPseq1 as part of the BRAIN Initiative BICCN project.^20^ Here we sought to replicate this MAPseq experiment with our updated protocol. We injected barcoded MAPseq virus into the primary motor cortex (Fig. 2a) and dissected 26 target regions matching the published MAPseq1 experiment,^20^ including regions in the cortex, striatum, thalamus, midbrain, and hindbrain. We then processed the dissected samples using the MAPseq2 protocol, yielding the projections of 11,153 neurons total in two mice. We clustered the published MAPseq1 data and our MAPseq2 data using agglomerative hierarchical clustering and identified the 5 expected cortical projection cell types^20^: intratelencephalic (IT) neurons with or without striatal or contralateral projections (ITi, ITc STR+, ITc STR–), corticothalamic (CT), and extratelencephalic (ET) neurons (Fig. 2b, c). We then matched the cell types discovered in MAPseq1 and MAPseq2 data to each other using using Pearson’s correlation or MetaNeighbor^35^ (Fig. 2d,e). As expected, all cell types matched with their appropriate partner. To further validate our MAPseq2 tracing data, we downsampled the MAPseq2 data to equal the number of neurons recovered in the published MAPseq1 data and combined the two datasets without any batch correction. After clustering the combined matrix, we again recovered all expected cell types. Importantly, MAPseq1 and MAPseq2 neurons were well intermingled within each cluster (Fig. 2f), and the proportion of cell types in MAPseq1 and MAPseq2 were comparable (Fig. 2g, h). Overall, this head-to-head comparison of MAPseq1 and MAPseq2 validates MAPseq2 in producing reliable single-cell projection data.

**Figure 2.**
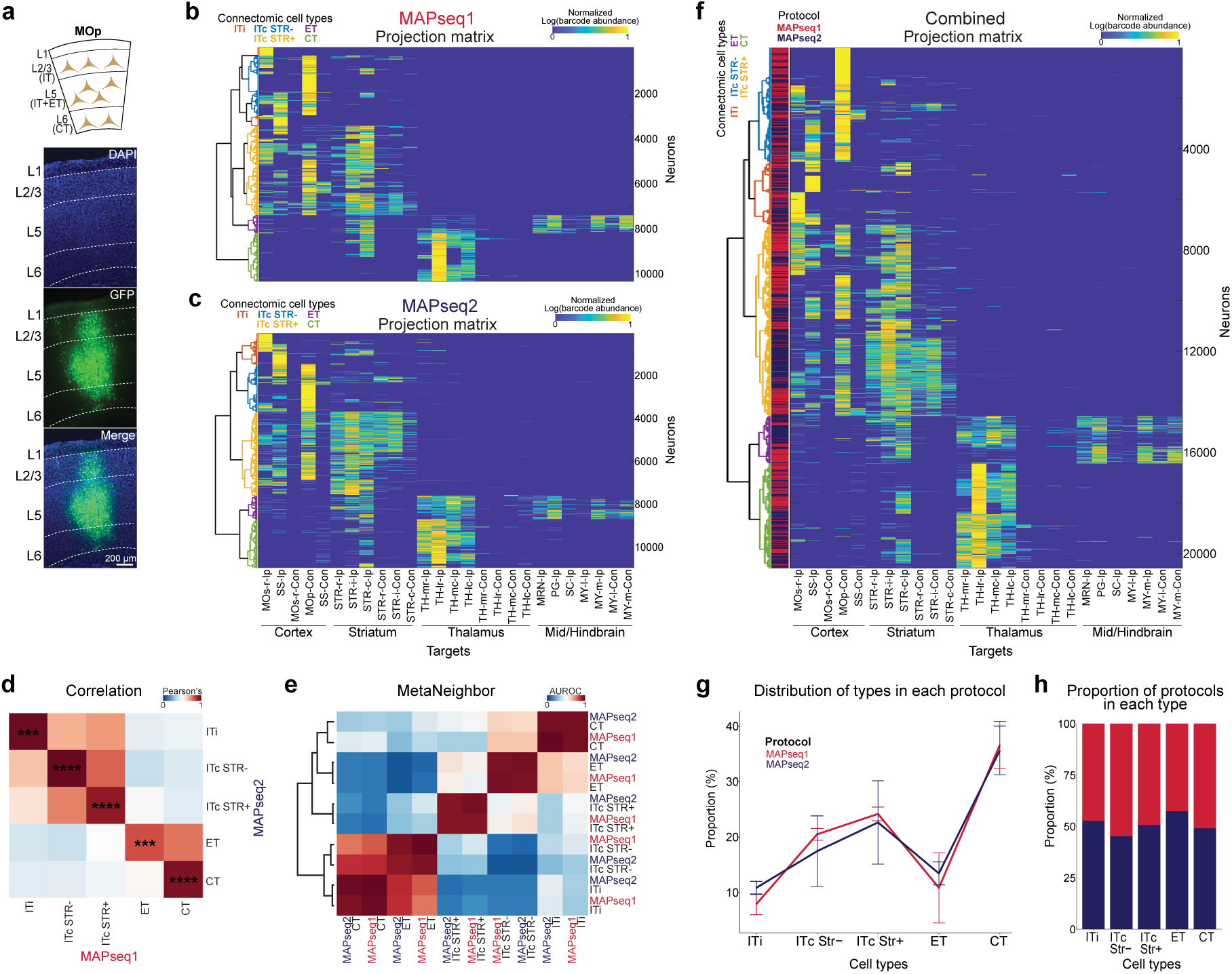
Validation of MAPseq2 for single-cell projection mapping. **a,** Layered structure of mouse primary motor cortex with connectomic cell types in each layer (top). Our injections of MAPseq virus cover all layers (bottom). **b, c,** Projection matrices of the mouse primary motor cortical neurons reconstructed by MAPseq1 (**b,** published MAPseq1 dataset,^19^ 2 mice, 10,295 neurons) and MAPseq2 (**c,** 2 mice, 11,153 neurons) with hierarchical clustering. Projections were mapped across 26 brain regions including forebrain, midbrain, and hindbrain regions. The parula color bar represents normalized barcode abundance, which was first normalized by the total barcode abundance across all targets within each neuron, log-transformed, and then normalized by the maximum value of the matrix bringing the scale as from 0 to 1. **d,** Correlation of connectomic cell types between MAPseq1 and MAPseq2. Pearson correlation was applied to the averaged projection strengths of all neurons in each type. Significance was determined by 10,000 permutations (*p < 0.05, **p < 0.01, ***p < 0.001, ****p < 0.0001). **e,** MetaNeighbor analysis of the connectomic cell types identified by MAPseq1 and MAPseq2. Higher AUROC score indicates greater similarity between the types. **f,** Combined projection matrix of MAPseq1 and MAPseq2 datasets. The MAPseq1 dataset of **b** was combined with a randomly subsampled set of MAPseq2 neurons from panel **c**, matching the number of neurons of the MAPseq1 dataset. Hierarchical clustering was then performed on the combined dataset. The protocol color bar shows good intermingling of MAPseq1 and MAPseq2 neurons across the connectomic cell types. **g, h** Comparable proportion of connectomic cell types between MAPseq1 and MAPseq2 in the combined projection matrix.

In conclusion, we here present and validate MAPseq2, which brings improved barcode detection sensitivity, reduced costs, and simplified procedures to barcoded connectomics techniques, including MAPseq, BRICseq, BARseq, ConnectID, and POINTseq.

## Methods

### Animals

We conducted all mouse procedures in accordance with the Johns Hopkins University Animal Care and Use Committee (ACUC) protocols MO20M376 and MO23M346. Mice were maintained on a 12-hour lightIdark cycle with ad libitum access to food and water. We purchased C57BLI6J mice from the Jackson Laboratory and maintained them in the lab. We used male offspring for experiments.

### MAPseq virus

To generate the MAPseq viral library, we first constructed a barcoded plasmid diversity library encoding the genomic MAPseq RNA, using the HZ120 plasmid kindly provided by Anthony M. Zador^32^ as described previously.^12^ The plasmid library had a diversity of 7 x 10^6^. We then produced a MAPseq viral library and assessed its titer as previously described.^12^ Briefly, we linearized the library plasmid and a Sindbis virus helper plasmid (Addgene #72309) with Pac I and Xho I, respectively, and synthesized RNA in vitro using the mMessage mMachine SP6 Transcription Kit (Invitrogen, AM1340). We transfected the synthesized RNAs into BHK cells at 80–90% confluence in 10 cm dishes using Lipofectamine 2000 (Invitrogen, 11668027) at a 1:1 molar ratio. BHK cells were cultured in a medium composed of MEM Alpha (Gibco, 12571063), 5% FBS (Cytiva, SH30070.03), 1x Antibiotic-Antimycotic (Gibco, 15240096), 1x MEM Vitamin Solution (Gibco, 11120052), and 1x L-Glutamine (Gibco, A2916801), maintained at 37°C with 5% CO_2_ in a humidified incubator under sterile conditions. We collected the culture medium 40–44 hours post-transfection, and concentrated the virus by ultracentrifugation at 160,000 × *g* and 4°C for 2 hours. The resulting viral library had a titer of 4.0 × 10^11^ vgImL determined by qPCR using a StepOnePlus system (Applied Biosystems, 4376600) after RT with a gene specific primer. The primer sequences were as follows.

RT primer: 5’-GGG TCG CCT TGC TTG AAG TG-3’

qPCR forward primer: 5’ TAT CCG CAG TGC GGT TCC AT 3’

qPCR reverse primer: 5’ TGT CGC TGA GTC CAG TGT TGG 3’

To examine the diversity of the viral library, we extracted viral genomic RNA from 4 x 10^8^ viral particles, produced a sequencing library by MAPseq2, and sequenced it, yielding > 1.8 × 10^6^ unique barcodes (Supplementary Fig. 6a). We then estimated the fraction of uniquely labeled neurons in each MAPseq experiment (F) using

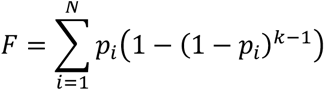

where k is the number of infected neurons, N is the number of unique barcodes (i.e., barcode diversity), and *p_i_* is the probability of each barcode based on the barcode distribution (Supplementary Fig. 6b).^12^ To avoid overestimating the diversity of the viral library, we only considered barcodes that were detected with at least 2 UMI counts when sequencing the virus. We found that across all experiments, the fraction of uniquely labelled neurons was close to 1, with the experiment with the highest number of neurons infected (MAPseq2-1: 7,506 neurons) yielding a fraction of 0.9904.

### Optimizing PCR1 and RT primers

To make PCR1 primers compatible with our new PCR2 primer designs, we used a new PCR1 reverse primer (5’ GAC GTG TGC TCT TCC GAT CT 3’) and redesigned the RT primer to be compatible to the updated PCR1 reverse primer (v1: 5’ GAC GTG TGC TCT TCC GAT CTN NNN NNN NNN NNX XXX XXX XTG TAC AGC TAG CGG TGG TCG 3’). N_12_ is the UMI, X_8_ the SSI sequence. We then tested four different forward primer sequences (A-D) to minimize non-specific amplification and chose primer D. The sequences are as follows:

A (used in MAPseq1)^12^: 5’ ACG AGC TGT ACA AGT AAA CGC GT 3’

B: 5’ CTG TAC AAG TAA ACG CGT AAT G 3’

C: 5’ TGC TGC CCG ACA ACC ACT AC 3’

D: 5’ CGA GAA GCG CGA TCA CAT G 3’

To further improve specificity, we redesigned RT primers with a different binding site (v2: 5’ GAC GTG TGC TCT TCC GAT CTN NNN NNN NNN NNX XXX XXX XCA CGA CGG CAA TTA GGT AGC 3’; N12 is the UMI, X8 the SSI sequence) and tested the v2 RT primers by performing RT and PCR1.

### Surgical procedures

For viral injections, we anesthetized mice with isoflurane (4% for induction, 1–1.5% for maintenance) and positioned them in a stereotaxic frame. We administered Lidocaine (5 mgIkg) and meloxicam (2 mgIkg) subcutaneously, made a midline scalp incision, and drilled a small craniotomy at the target coordinates. We then injected MAPseq virus (4 x 10^11^ vgIml) using pulled glass micropipettes and a Nanoject III injector (Drummond Scientific) at a rate of 1 nLIsec, closed the surgery site, and cared for the animals postop according to institutional ACUC guidelines.

For comparison of MAPseq1 and MAPseq2, we injected 200 nL of virus at each of 3 depths in the left MOp (AP: +0.5 mm, ML: +1.5 mm, DV: -0.3, -0.6, -1.0 mm relative to bregma). For comparison of SUPERSCRIPT IV and MashUp, we injected 200 nL of virus in the barrel cortex (AP: -1.2 mm, ML: +3.5 mm, DV: -0.6 mm relative to bregma).

### Imaging

40-44 hours after MAPseq virus injections, we euthanized the mice, transcardially perfused then with PBS and 4% paraformaldehyde (PFA), and collected their brains, followed by a 24-hour fixation in 4% PFA at 4°C. After fixation, we stored the brains in 30% sucrose at 4°C for 24 hours and froze them at -80°C with OCT. We cryosectioned the fixed brains coronally at 50 µm thickness and mounted them on slides with Fluoromount-G Mounting Medium with DAPI (Invitrogen, 00-4959-52). Images of the slices were taken using a fluorescence microscope (Keyence, BZ- X710) using a 4x objective (NA = 0.20).

### Generation of spike-in RNA

We generated spike-in RNAs for sample normalization in MAPseq^12^ by in vitro transcribing RNA from a MAPseq plasmid library (diversity 3x10^6^) using the mMessage mMachine SP6 Transcription Kit (Invitrogen, AM1340). To differentiate spike-in RNA from viral RNA, we designed the barcode region to contain a 24 nt random sequence followed by a constant 8 nt identifier (CGTCAGTC). During library preparation, we added 10^5^ spike-in molecules to each target sample and 10^5^ or 10^7^ molecules to each source sample prior to reverse transcription.

### MAPseq2 procedures

40-44 hours after MAPseq virus injections, we euthanized the mice and collected their brains. We immediately snap froze the fresh brains at -80°C with OCT (Electron Microscopy Sciences). We cryosectioned the fresh frozen brains coronally at 300 µm thickness and mounted the sections on glass slides. We then manually dissected the target regions by placing the slides on a metal plate surrounded by 2.25 M CaCl_2_, which was continuously chilled with dry ice to maintain a temperature of approximately -20 °C.^36^ After dissection, we added TRIzol (Invitrogen, 15596026) to each sample and homogenized the tissue with pestles (Fisherbrand, 12-141-368), followed by RNA extraction according to the manufacturer’s instructions.

After the TRIzol RNA extraction, we performed RT using the custom MashUp RT enzyme (https://pipettejockey.com/), produced for us by UCBerkely QB3 Macrolab, with sample-specific RT primers and spike-in RNA. We followed the MashUp RT protocol provided by its inventor (https://pipettejockey.com/) with modifications. Briefly, we diluted 1x MashUp enzyme as supplied by the Marcolab in 1x storage buffer (150 mM NaCl, 50 mM Tris-HCl pH 7.5, 1 mM EDTA, 0.1% IGEPAL CA-630, 50% glycerol) to 0.02x (50 times dilution) as this was the optimal MashUp concentration for RT in MAPseq2. We performed MashUP RT similarly to SuperScript IV RT. Briefly, we incubated RNA, dNTP, and RT primer solution at 70°C for 10 mins and cooled it down on ice. Then we added DTT, RNase inhibitor, MashUp enzyme, and 5x MashUp buffer (125 mM Tris-HCl pH 8.3, 125 mM MOPs pH 7.9, 300 mM KCl, 20 mM MgCl_2_, 25% glycerol, 0.03% NP-40) and incubated the mix at 55°C for 1 hour. We then cooled it down to 37°C and added Exo I (NEB, M0568L) in the tube and incubated for 15 mins and inactivated MashUP and Exo I at 80°C for 10 mins. We then cooled it down to 37 °C, added RNase If (NEB, M0243L), and incubated for 1 hour, followed by 20 mins incubation at 80 °C to inactivate RNase If. The RT primers have the following sequence, 5’ GAC GTG TGC TCT TCC GAT CTN NNN NNN NNN NNX XXX XXX XCA CGA CGG CAA TTA GGT AGC 3’, where N_12_ corresponds to the UMI and X_8_ corresponds to the predetermined SSI. Then, we performed a bead clean-up of the RT products using 1.8X home-made or AMPure XP (Beckman Coulter, A63881) beads according to the manufacturer’s instructions. For qPCR of PCR1, we prepared 5x qPCR master mix using AccuPrime Pfx polymerase (Invitrogen, 12344024)^12^. The master mix consisted of 5x SYBR green I (Invitrogen, S7563), 5x AccuPrime reaction mix, 2.5mM MgSO_4_, 5µM foward primer, 5µM reverse primer, 10% ROX Reference Dye (Invitrogen,12223012), and 25% DMSO. We added the qPCR master mix to the cleaned RT product and ran qPCR using StepOnePlus (Applied Biosystems, 4376600) or LightCycler96 (Roche, 05815916001), monitoring the amplification in real-time and stopped the reaction before reaching saturation. The forward and reverse primers of PCR1 have following sequences, 5’ CGA GAA GCG CGA TCA CAT G 3’ and 5’ CTG GAG TTC AGA CGT GTG CTC TTC CGA TCT 3’, respectively. We performed PCR2 in the same way using diluted PCR1 products as templates except that sample-specific primers with specific i5 and i7 were used. The forward and reverse primers of PCR2 have following sequences, 5’ AAT GAT ACG GCG ACC ACC GAG ATC TAC ACX XXX XXX XAC ACT CTT TCC CTA CAC GAC GCT 3’ and 5’ CAA GCA GAA GAC GGC ATA CGA GAT XXX XXX XXG TGA CTG GAG TTC AGA CGT GTG CTC TTC 3’, respectively. The X_8_ of the forward and reverse primers correspond to i5 and i7 indexes, respectively. We then extracted the expected 219bp PCR2 band using QIAquick gel extraction kit (Qiagen, 28704) following the manufacturer’s instructions. We measured the size and concentration of the extracted products with a TapeStation (Agilent, G2992AA). For sequencing, we submitted our MAPseq2 libraries to a sequencing core at Johns Hopkins University. The libraries were sequenced on NovaSeq X Plus platforms, resulting in paired-end reads of R1 and R2, demultiplexed based on the i5 and i7 pairs. R1 was at least 32 nt in length starting from the barcode region, unless otherwise specified. R2 was at least 20 nt long, covering SSI and UMI.

### Assessment of Exo I treatment efficiency and the first-strand cDNA protection from Exo I

To evaluate the efficiency of Exo I treatment and the protection of first-strand cDNA, we infected BHK cells with MAPseq virus (1 x 10^7^ vgIwell) in 24-well plates and incubated them for 24 hours. We collected the cells, homogenized them with pestles, and extracted total RNA using TRIzol reagent (Invitrogen, 15596026) according to the manufacturer’s protocol. We then performed RT following the MAPseq2 protocol, with or without Exo I treatment. To assess the presence of residual RT primers in the RT products, we ran samples on a 15% TBE-Urea gel (Invitrogen, EC6885BOX) under denaturing conditions and visualized ssDNA using SYBR Green II RNA gel stain (Invitrogen, S7564). To assess the protection of the first-strand cDNA, we conducted qPCR to measure the amount of barcode cDNA using a StepOnePlus system (Applied Biosystems, 4376600) and HotStart™ 2X Green qPCR Master Mix (Apexbio, K1070). The following forward and reverse primers were used, 5’ GAC GAC GGC AAC TAC AAG AC 3’ and 5’ TAG TTG TAC TCC AGC TTG TGC 3’.

### Assessment of tolerable background RNA range

To evaluate how varying amounts of endogenous total RNA affect MAPseq2 performance, we added 10^5^ molecules of spike-in RNA to 0.1 µg, 1 µg, 5 µg, or 10 µg of total RNA extracted from a mouse brain, and performed MAPseq2 RT, ExoI and RNAse If treatment, followed by bead cleanup, and PCR1. MAPseq2 PCR1 amplification efficiency is defined as the fold-change of dsDNA fluorescence per PCR cycle (ranging from 1 to 2). To measure this, we identified the exponential phase of the amplification plot by selecting the most linear segment of four consecutive points below the Cq value on the log transformed plot and calculated the slope.

### Home-made TRIzol and beads

We prepared home-made TRIzol as described previously,^27,28^ consisting of 38% phenol-chloroform, 0.8M guanidine thiocyanate, ammonium thiocyanate, 0.1M sodium acetate, and 5% glycerol. For the optimized reagent, we increased the guanidine thiocyanate concentration to 1.7M. We also produced home-made beads as previously described.^31^ Briefly, we washed Sera-Mag SpeedBeads (Fisher Scientific, Cat.# 09-981-124) with TE buffer and then resuspended the beads in a solution consisting of 18% PEG-8000, 1M NaCl, 10mM Tris-HCl, 1mM EDTA, and 0.055% Tween-20 as 2% of the total volume. Both reagents substitute 1:1 for the commercially available reagents in standard protocols.

To test the performance of home-made TRIzol, we extracted RNA from mouse cortical tissue punches of the same size, following the manufacturer’s instructions for commercial TRIzol. We performed RT using MashUp and an oligo- dT primer, and quantified the housekeeping gene Actb, which encodes β-actin, using qPCR on the StepOnePlus system (Applied Biosystems, 4376600) and the following primers:

Actb forward: 5’-CGG TTC CGA TGC CCT GAG GCT CTT-3’

Actb reverse: 5’-CGT CAC ACT TCA TGA TGG AAT TGA-3’

To test the performance of home-made beads, we conducted MAPseq2 procedures up to the clean-up step, using 10^5^, 10^6^, and 10^7^ molecules of amount of spike-in RNA. We then performed clean-up using either AmpureXP or home-made beads, followed by PCR1. The Cq value and amplification efficiency were then measured.

### Producing raw barcode matrices from MAPseq2 sequencing results

The procedure is based on the original MAPseq sequencing data processing pipeline.^12^ Briefly, after demultiplexing by i5 and i7 indexes, each pair of read1 and read2 FASTQ files corresponded to a specific brain region, and we processed the files using bash. First, we combined the first 32 nt of read1 and the first 20 nt of read2 and filtered based on the SSI information. Since we used a specific SSI, i5, and i7 for each sample, most of the demultiplexed reads should contain the correct SSI, and we found that 98-99% of our reads had the correct SSI. Next, we removed the SSI sequence, and used the remaining 44 nt sequences to generate rank plots of each sequence and its abundance using MATLAB (MathWorks, R2022a). Rank plots exhibit a plateau and a tail, where the plateau includes reliable sequences and the tail includes primarily sequencing and PCR errors and template switches.^24^ We manually determined minimal read thresholds for all samples to exclude the tail and collapsed the filtered sequences into unique 32 nt sequences now comprising only the 30 nt barcode and 2 nt pyrimidine anchor and measured their abundance as the number of different UMIs each 32mer was seen with. We then split viral barcodes from spike-in barcodes, based on whether they contained the spike-in specific sequence (CGTCAGTC). To correct for sequencing errors, we collapsed all 32 nt sequences within a Hamming distance of 3 or less to the most abundant barcode sequence in each group. To achieve this, we constructed a connectivity matrix of all 32 nt sequences using MATLAB, where connections were determined by Bowtie alignments of the barcodes^37^ allowing up to 3 mismatches. In each connected component of the matrix, the most abundant sequence represented the group. Finally, we matched barcodes across samples and constructed a raw barcode matrix (barcode x region) as a MATLAB variable, where each element represented the number of UMIs of the barcode.

### Producing projection matrices

We analyzed barcode matrices in MATLAB. We filtered barcodes to include only high quality neurons. To do so, we only included barcodes that have more than 10 UMIs at their strongest target region. Additionally, to consider only well barcoded cells, we set a UMI threshold in the source region. To determine an optimal threshold, we plotted two parameters: 1) the number of filtered barcodes, and 2) the proportion of barcodes projecting to only one region. Based on the results, we set the source threshold at 20 UMIs, where these parameters stabilize. Then, we normalized barcode abundances in each region by the amount of spike-in RNA in that region. We further normalized barcode abundances across regions for each barcode by the sum of that barcode across all target regions. Finally, we log transformed all values and scaled them so that the maximal projection value was 1.

### Assessment of MAPseq2 barcode recovery efficiency

To estimate the performance of MAPseq2 compared to MAPseq1 using same samples, we injected MAPseq virus (4 x 10^11^ vgIml, 200 nl) in the primary motor cortex of the left hemisphere (AP: +0.5 mm, ML: +1.5 mm, DV: -0.6 mm relative to bregma). 40-44 hours after the injection, we collected the brain and extracted total RNA from the injection site. We then diluted the sample 100 times or 10 times with RNase-free water to generate multiple replicate samples. For MAPseq1, we sent the samples to the MAPseqIBARseq core at Cold Spring Harbor Laboratory. For MAPseq2, we processed the samples and sequenced them according to the MAPseq2 protocol, with a Read1 length to 28nt. We created raw barcode matrices from the 100 times or 10 times diluted samples of MAPseq1 and MAPseq2 (4 groups: MAPseq1 with 100 times diluted, MAPseq2 with 100 times diluted, MAPseq1 with 10 times diluted, and MAPseq2 with 10 times diluted). The mean value of the 3 replicates from each group was used as the UMI count for each barcode. Afterward, we performed linear regression analysis using R (version 4.4.1).

We also compared SuperScript IV and MashUp RT enzyme using an RNA sample extracted from a MAPseq virus- injected mouse barrel cortex tissue. The undiluted RNA from the injection site was divided equally into two tubes, and each was processed with either SuperScript IV or MashUp. For the SuperScript IV condition, we performed the RT as described for MashUp above, except that we used the SuperScript IV buffer and incubated for 10 minutes for the reaction after adding enzyme and buffers according to the manufacturer’s protocol, whereas the MashUp RT was incubated for 1 hour. Barcode recovery efficiency was analyzed using the same pipeline described above but using 32nt of Read 1.

### Assessment of MAPseq2 for projection mapping

We first obtained MAPseq1 data files from a published study.^20^ Specifically, we used the “projnorm” variable from “M1BARseqdata.mat”, which represents projection matrices of 4,729 and 5,570 neurons from two mice across 40 targets. We reduced the number of targets to 26, aligning them with our dissected target regions. Rows with 0 UMI counts across all 26 regions were then removed, resulting in a MAPseq1 projection matrix consisting of 10,295 neurons.

We then performed hierarchical clustering on the rows/barcodes of the MAPseq1 projection matrix using a Ward’s method with an Euclidean distance metric, using MATLAB’s “linkage” and “dendrogram” functions. To identify major clusters, we set the “maxclust” parameter of the dendrogram function to a specific number, which cuts the dendrogram at a position that yields that number of clusters. After unbiased clustering, we sorted the rows according to the order of clusters in the dendrogram and plotted the matrix. To identify the 5 major types of MAPseq1 dataset, ITi, ITc STR-, ITc STR+, CT, and ET,^20^ we cut the dendrogram ending up with 8 clusters (ITi 1, ITc STR- 1, ITc STR- 2, ITi 2, ITc STR+ 1, ITc STR+ 2, ET, and CT). Based on the projection patterns of ITi, ITc STR-, ITc STR+, ET, and CT, we merged the clusters yielding 5 final clusters.

For MAPseq2, we injected MAPseq virus in MOp and 40-44 hours after the injection, we collected the brains for imaging or MAPseq analysis. We dissected the source, MOp, and 26 target regions (Data S1) and processed the samples following the MAPseq2 protocol. We processed two mice, MAPseq2-1 and MAPseq2-2, obtaining projection matrices as described above, which contain 7,506 and 3,464 neurons with 26 target regions, respectively. We then performed hierarchical clustering on the MAPseq2 dataset by setting the number of clusters to 5. Based on their projection patterns, we assigned the resulting clusters to ITi, ITc STR–, ITc STR+, ET, and CT.

To assess the similarity between projection types identified by each protocol, we calculated the mean projection values for each type and computed pairwise Pearson correlations between types using R. In parallel, we conducted MetaNeighbor^35^ analysis between the types identified by each protocol, where higher AUROC scores indicate higher similarity between the two types.

To compare MAPseq1 and MAPseq2 based on clusters generated under the same criteria, we randomly selected 10,295 neurons from the MAPseq2 dataset to match the size of the MAPseq1 dataset and combined them into a joint projection matrix (20,590 neurons x 26 target regions). We performed hierarchical clustering on the combined dataset by setting the number of clusters to 5. Based on their projection patterns, we assigned the resulting clusters to ITi, ITc STR–, ITc STR+, ET, and CT.

### Statistical analyses

We performed student’s t test, one-way ANOVA with Bonferroni post-hoc tests, linear regression analysis, and Pearson correlation test using R functions (“t.test”, “aov”, “pairwise.t.test”, “lm”,”cor”). MetaNeighbor analysis was conducted using MetaNeighbor R package (version 1.25.0). We conducted all tests as two-sided, and their significance levels are noted in the text and figures.

## Data availability

Dissection images of brain regions are provided as Supplementary File 1. All raw sequencing datasets are available in the NCBI Sequence Read Archive (SRA) under the following BioSample accession numbers: MAPseq virus library (SAMN53393094), barrel cortex (SAMN49783975), motor cortex diluted replicates (SAMN49783974), MAPseq2-1 (SAMN53373349), and MAPseq2-2 (SAMN53373327). All processed MAPseq data are available as Supplementary File 2.

## Code availability

All analysis scripts used for MAPseq data processing is available on GitHub (https://github.com/HyopilKim/MAPseq2-and-POINTseq).

## Acknowledgements

We thank Anthony M. Zador for generously providing us with plasmid construct HZ120.^32^ We also appreciate insightful discussions on MAPseq1 and MAPseq2 with Huiqing Zhan and Xiaoyin Chen. This work was supported by a SFARI pilot award 875575, and NIH grants U01NS132161, DP1DA056668, and RF1AG078378 to J.M.K. and a postdoctoral fellowship from the Autism Science Foundation to H.K.

## Author information

### Contributions

H.K. and J.M.K conceptualized the study and prepared the manuscript. H.Q. assessed the limit of background RNA input for MAPseq2. H.K. and C.W. generated sequencing libraries. C.W. prepared the home-made beads. X.L. prepared the home-made TRIzol substitute. H.K. performed all the other experiments and analyses.

### Corresponding author

Correspondence to Justus M. Kebschull

## Ethics declarations

We declare no competing interests.

---

**Supplementary Figure 1.**
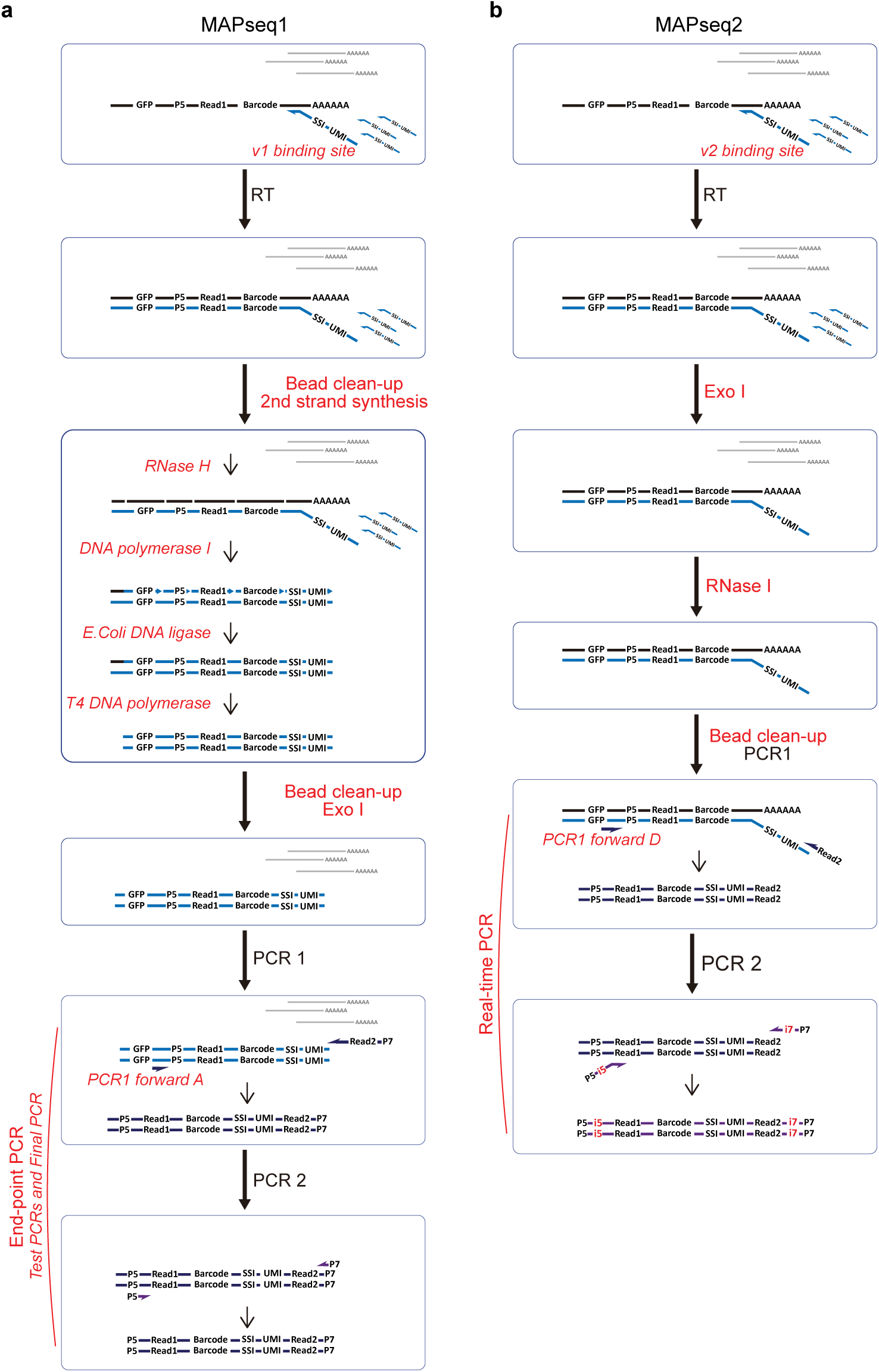
Detailed workflow of MAPseq2 compared to MAPseq1. **a, b,** Processes of sequencing library production by MAPseq1 (**a**) and MAPseq2 (**b**). Black line: viral RNA, Gray line: non-viral background RNA, Blue arrow: RT primers, Blue line: RT product, Navy arrow: PCR1 primer, Navy line: PCR1 product, Purple arrow: PCR2 primer, Purple line: PCR2 product. P5 and P7: adaptors for Illumina sequencing. i5 and i7: dual indexes for the patterned flow cell. Red indicates different materials or steps between MAPseq1 and MAPseq2.

**Supplementary Figure 2.**
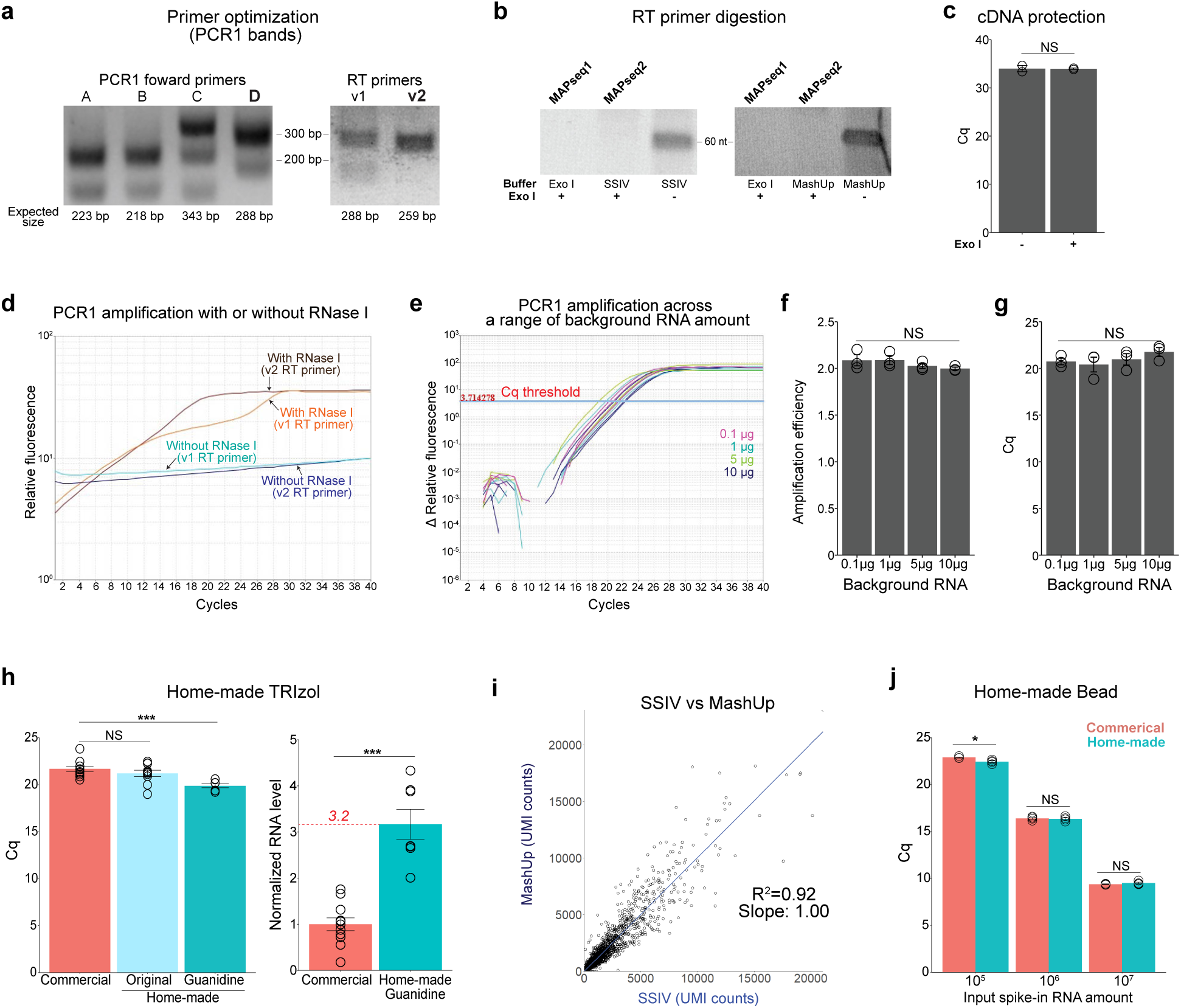
Optimization of MAPseq2. **a,** Optimization of PCR1 forward and RT primers. PCR1 forward primer D and v2 RT primer were selected as they produce the strongest target bands. For the PCR1 forward primer test, v1 RT primers were used and for RT primer test, PCR1 forward primer D was used. **b, c,** Evaluation of Exo I treatment on the first-strand cDNA. **b,** To digest RT primers (60 nt), in MAPseq1, Exo I is applied with Exo I buffer after the clean-up of the second-strand synthesis reaction, whereas in MAPseq2, Exo I is applied with SuperScript IV or MashUp buffer following the first-strand synthesis. Exo I activity for digesting RT primers in either SuperScript IV or MashUp RT buffers was examined and compared with its activity in the Exo I buffer. The successful digestion of RT primers is indicated by the absence of the 60 nt band on a denaturing PAGE gel. **c,** Exo I treatment on the first-strand cDNA after RT but before denaturation does not digest the first-strand cDNA (N=2 each). **d,** Representative samples with high total RNA input show reduced PCR1 amplification efficiency with both v1 and v2 RT primers. This was resolved by RNase I pretreatment. **e, f,** Influence of background RNA on PCR1 amplification. Comparable amplification plots (**e**) and amplification efficiency (fold-change per cycle) (**f**). MAPseq2 allows optimal barcode library preparation in the presence of a wide range of background RNA, at least up to 10 µg per sample, as measured by PCR1 amplification efficiency (N=3 each, 0.1–10 µg per sample; one-way ANOVA with Bonferroni post-hoc test, NS: not significant) **g**, Influence of background RNA on Cq values of PCR1. Comparable Cq values indicate consistent RT efficiency and cDNA yield across a wide range of background RNA in MAPseq2 (N=3 each). **h,** Comparison of total RNA extraction among commercial TRIzol (N=11), home-made substitute prepared as previously described^27,28^ (N=10), and optimized substitute with increased guanidine concentration (N=7). The recovered amount of *Actb* mRNA from fresh-frozen mouse brain tissues using the different reagents was measured by qRT-PCR. The optimized home- made TRIzol substitute achieved 3.2-fold higher RNA extraction (Student’s t test, NS: not significant, ***p < 0.001). **i,** Comparison of barcode recovery between a commonly used RT enzyme, SuperScript IV, and MashUp, performed on an injection site RNA sample (linear regression, R^2^=0.92, slope=1.00, *p*<0.0001). Each dot represents a barcode. **j**, Home-made beads performed similar to AmpureXP beads for RT product clean-up, which was assessed by Cq values of PCR1 (N=4 each, Student’s t test, NS: not significant).

**Supplementary Figure 3.**
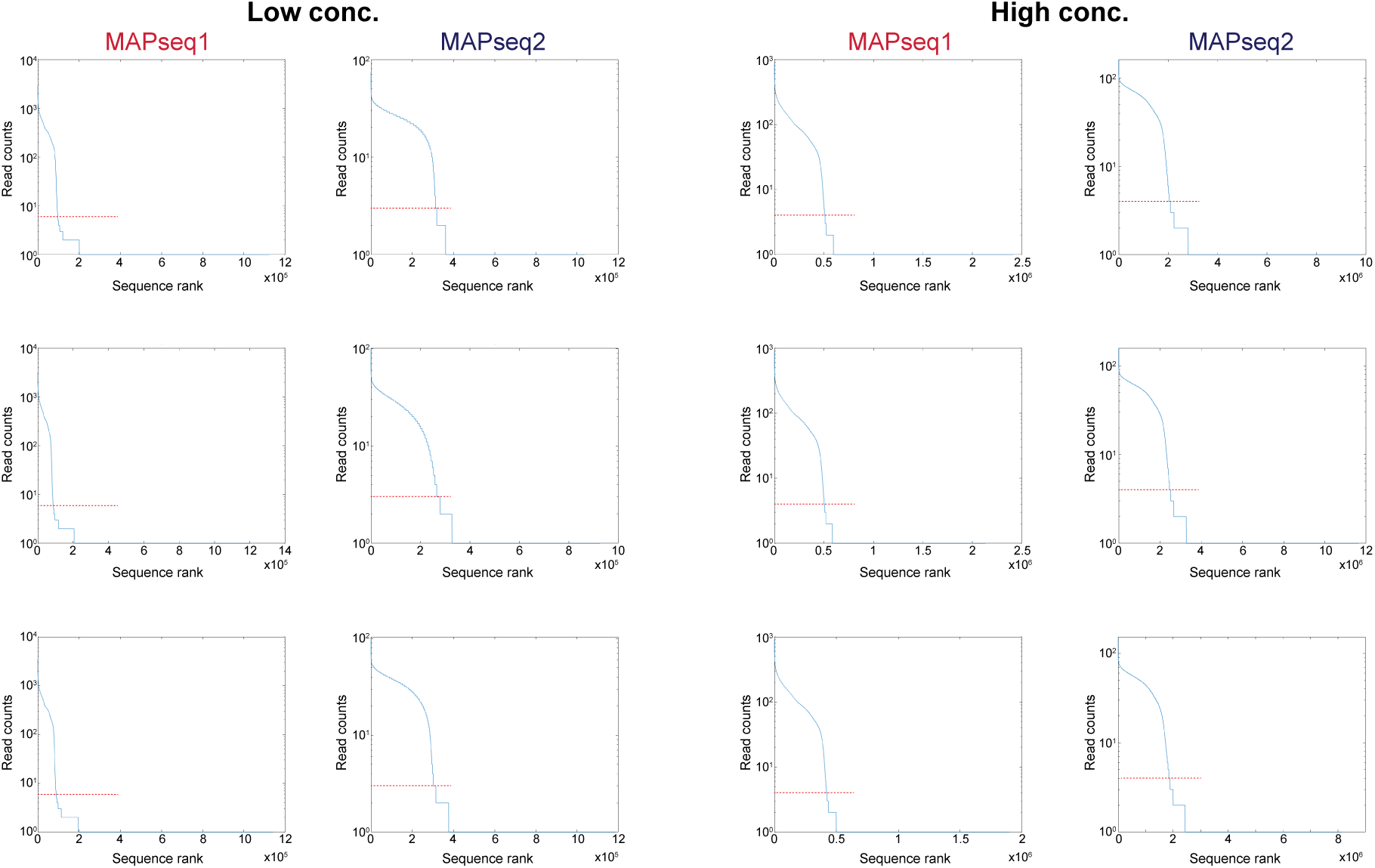
Rankplots of MAPseq1 and MAPseq2 data used in Figure 1. For each replicate sample, every unique barcode-UMI sequence is sorted along the x axis by read count, with the read counts plotted on the y axis. Red dashed lines indicate the thresholds used to filter for reliable reads.^12,24^ All samples were sequenced to saturation to avoid differences resulting from different sequencing depths between MAPseq1 and MAPseq2 samples.

**Supplementary Figure 4.**
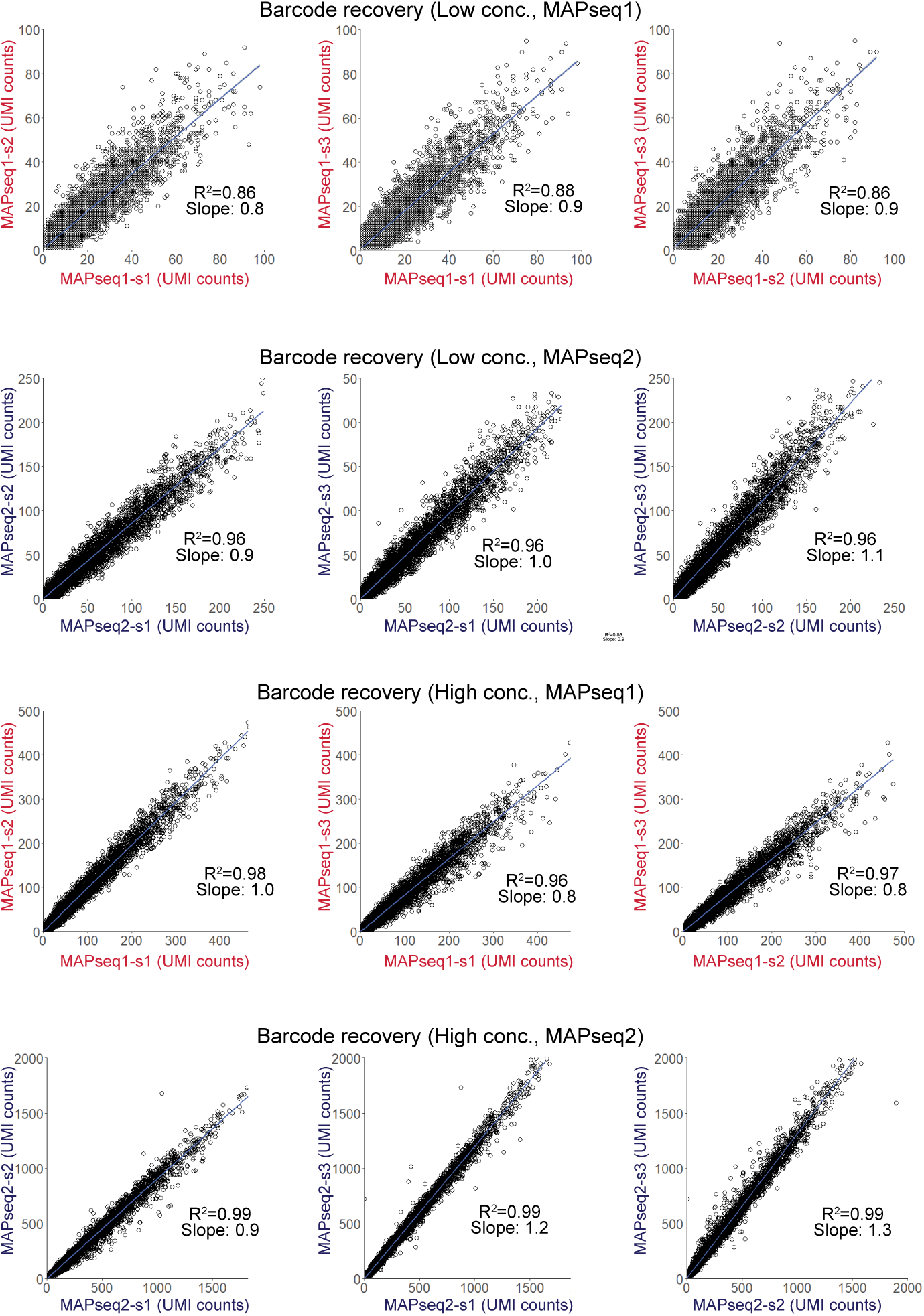
Correlation of barcode recovery between replicate samples. Barcode abundance is highly correlated between replicates within each of the four conditions (MAPseq1 low conc., MAPseq2 low conc., MAPseq1 high conc., and MAPseq2 high conc.). The 3 replicates of each condition were labeled as sample s1, s2, and s3.

**Supplementary Figure 5.**
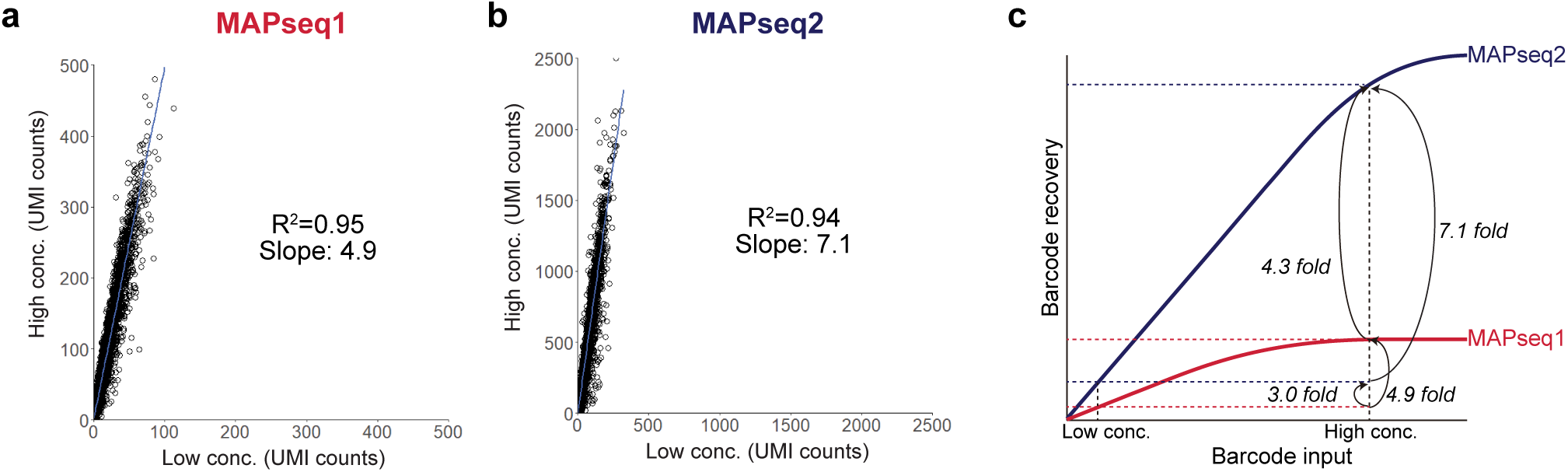
MAPseq2 has a higher capacity for barcode detection per sample than MAPseq1. **a, b,** Comparison of barcode recovery between the low (100 times diluted) and high (10 times diluted) concentration samples using the data from Fig. 1e, f. Each dot represents a barcode. Mean UMI values from three replicates are plotted (**a**, linear regression, R^2^=0.95, slope=4.9, *p*<0.0001; **b**, linear regression, R^2^=0.94, slope=7.1, *p*<0.0001). **c**, A model for barcode recovery of MAPseq1 and MAPseq2 according to the input barcode amount. In MAPseq, as in any other enzymatic method, barcode recovery presumably saturates when there are too many input barcodes. Saturation appears to occur earlier in MAPseq1 than in MAPseq2, resulting in the higher recovery fold change of MAPseq2 vs MAPseq1 for the high concentration samples compared to the low concentration samples.

**Associated Supplementary Note:**

In MAPseq1, source regions, which contain substantially more barcode mRNAs than target regions, generally show lower barcode recovery efficiency, suggesting an upper limit for how many barcode mRNA molecules can be detected in one MAPseq1 sample.^12^ Such an upper limit can result in undersampling of barcodes in high barcode samples. This effect is partially combated by normalizing barcode abundance per sample by an internal spike-in RNA, as is standard in MAPseq. However, severe undersampling can pose challenges in MAPseq injection sites, where undersampling may reduce the number of neurons that can be recovered, and in BRICseq, where both rare and abundant barcodes need to be detected in individual samples. The difference in performance of MAPseq2 vs. MAPseq1 in high vs. low barcode concentration samples indicates that MAPseq2 has a higher barcode detection capacity than MAPseq1 (Fig. 1e,f, Supplementary Fig. 5). To further explore this idea, we compared the amounts of barcodes recovered in the high vs. low concentration samples for each protocol. A priori, we expected a 10-fold difference in barcode abundances between the two concentrations. In MAPseq1, we observed only a 4.9-fold difference (Supplementary Fig. 5a). In contrast, MAPseq2, while not reaching the 10-fold difference either, showed a substantially improved fold change of 7.1 (Supplementary Fig. 5b), again indicating a higher capacity for MAPseq2 (Supplementary Fig. 5c).

**Supplementary Figure 5.**
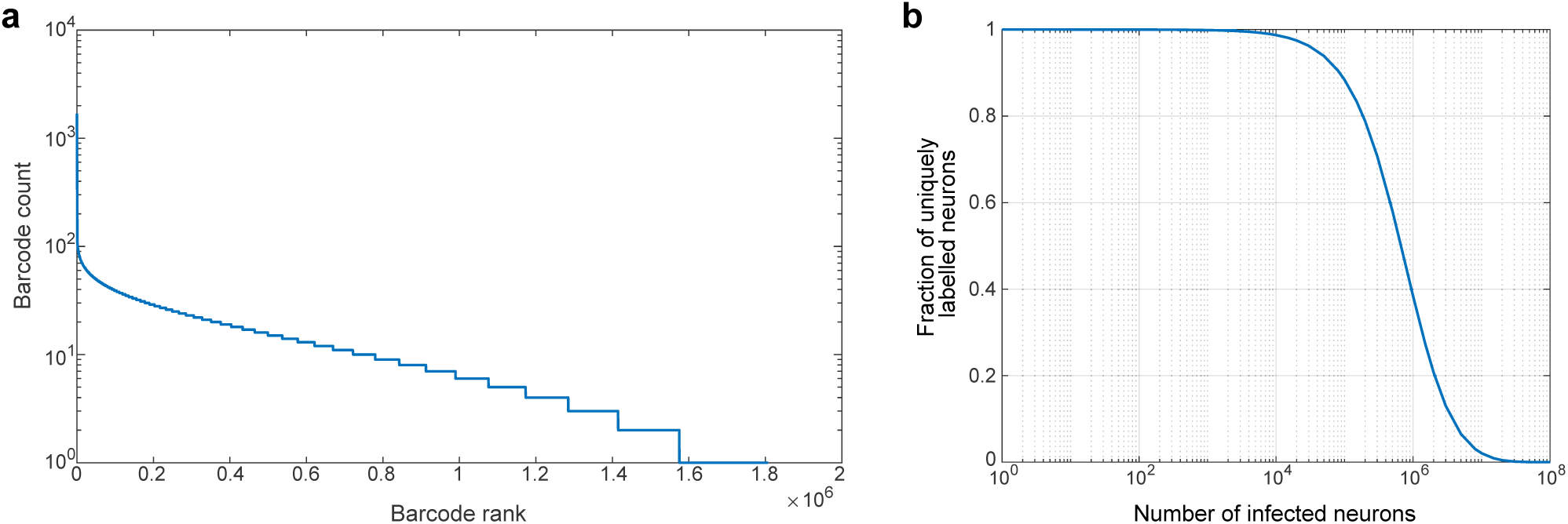
Diversity of virus used for MAPseq2. **a,** Rank plot of the viral barcode library by UMI counts that contains 1.6 × 10^6^ unique barcodes. **b,** Fraction of uniquely labeled neurons estimated from the rank plot in **a**, considering only barcodes with at least 3 UMI counts.

**Table S1.**
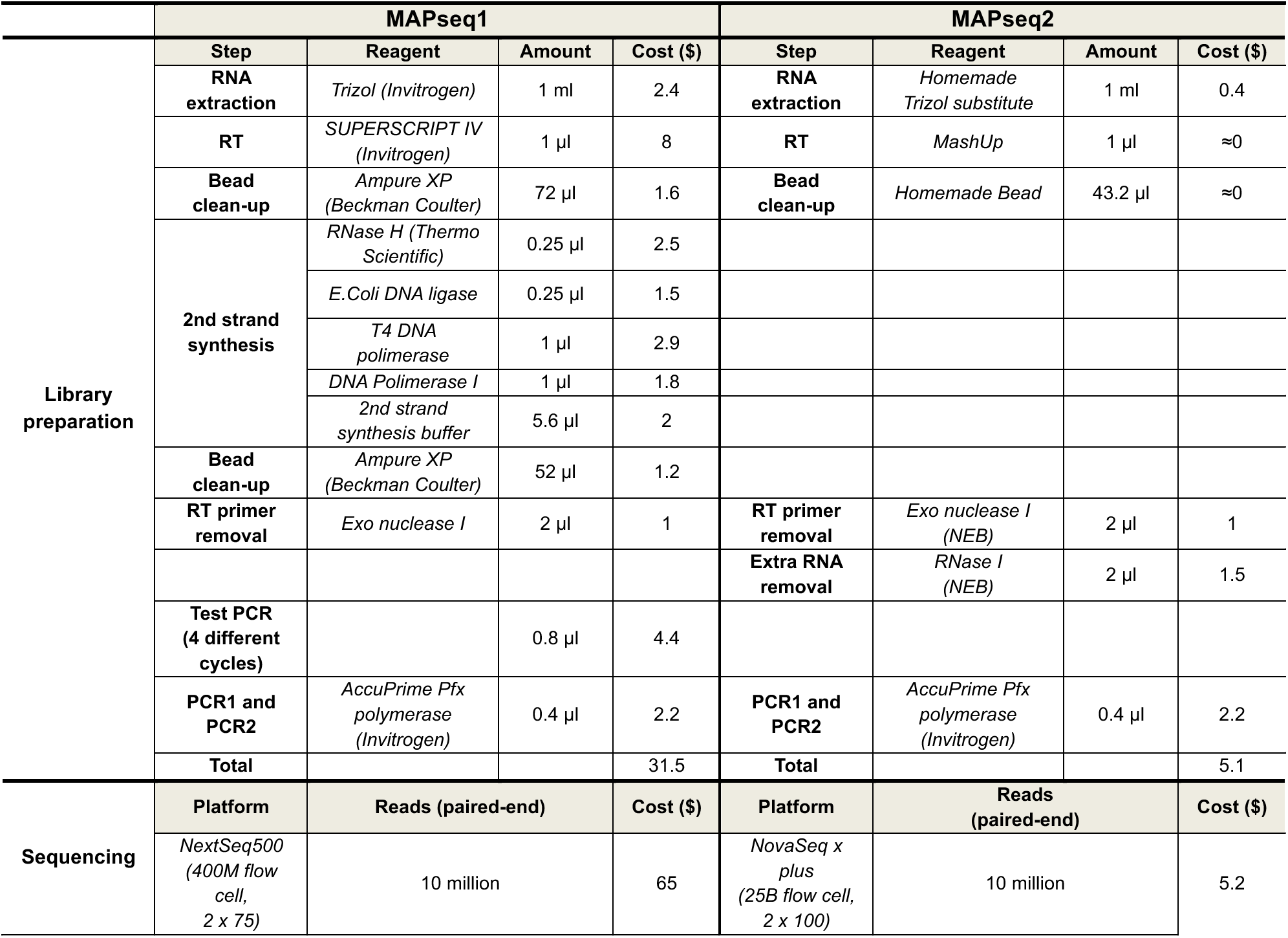
Per sample costs for MAPseq1 and MAPseq2. Costs less than $0.10 were not included. 10 million reads are typically appropriate for a target region. However, this sequencing depth may need to be adjusted for specific experiments.

## Notes

### Competing Interest Statement

The authors have declared no competing interest.

### Summary of Updates

we updated funding information to fully reflect the sources of funding listed in the paper.

