## Supplementary Data 1 for "MAPseq2: a sensitive barcoded connectomics method"

### Slide 1
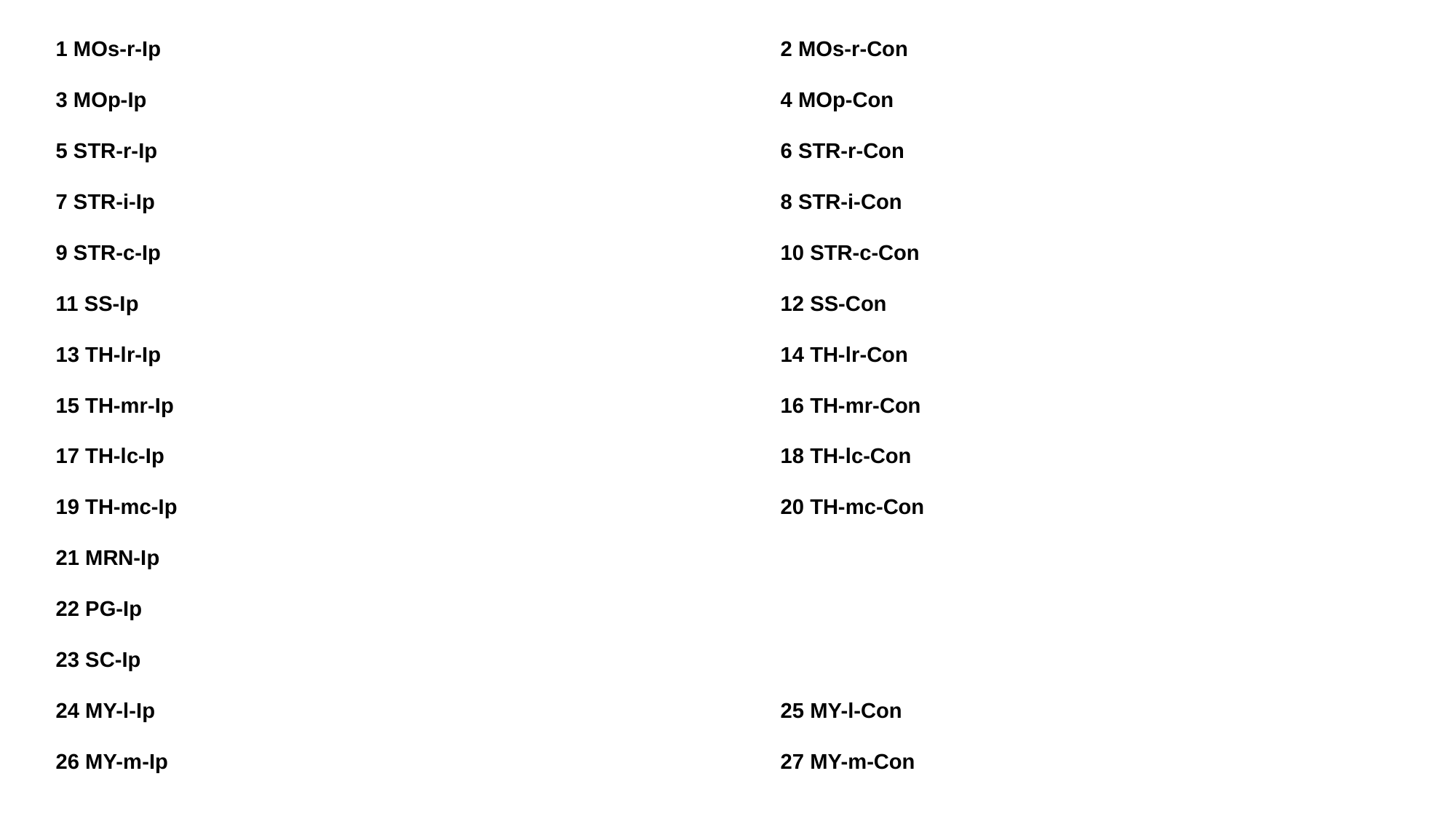

1 MOs-r-Ip
3 MOp-Ip
5 STR-r-Ip
7 STR-i-Ip
9 STR-c-Ip
11 SS-Ip
13 TH-lr-Ip
15 TH-mr-Ip
17 TH-lc-Ip
19 TH-mc-Ip
21 MRN-Ip
22 PG-Ip
23 SC-Ip
24 MY-l-Ip
26 MY-m-Ip
2 MOs-r-Con
4 MOp-Con
6 STR-r-Con
8 STR-i-Con
10 STR-c-Con
12 SS-Con
14 TH-lr-Con
16 TH-mr-Con
18 TH-lc-Con
20 TH-mc-Con
25 MY-l-Con
27 MY-m-Con

### Slide 2
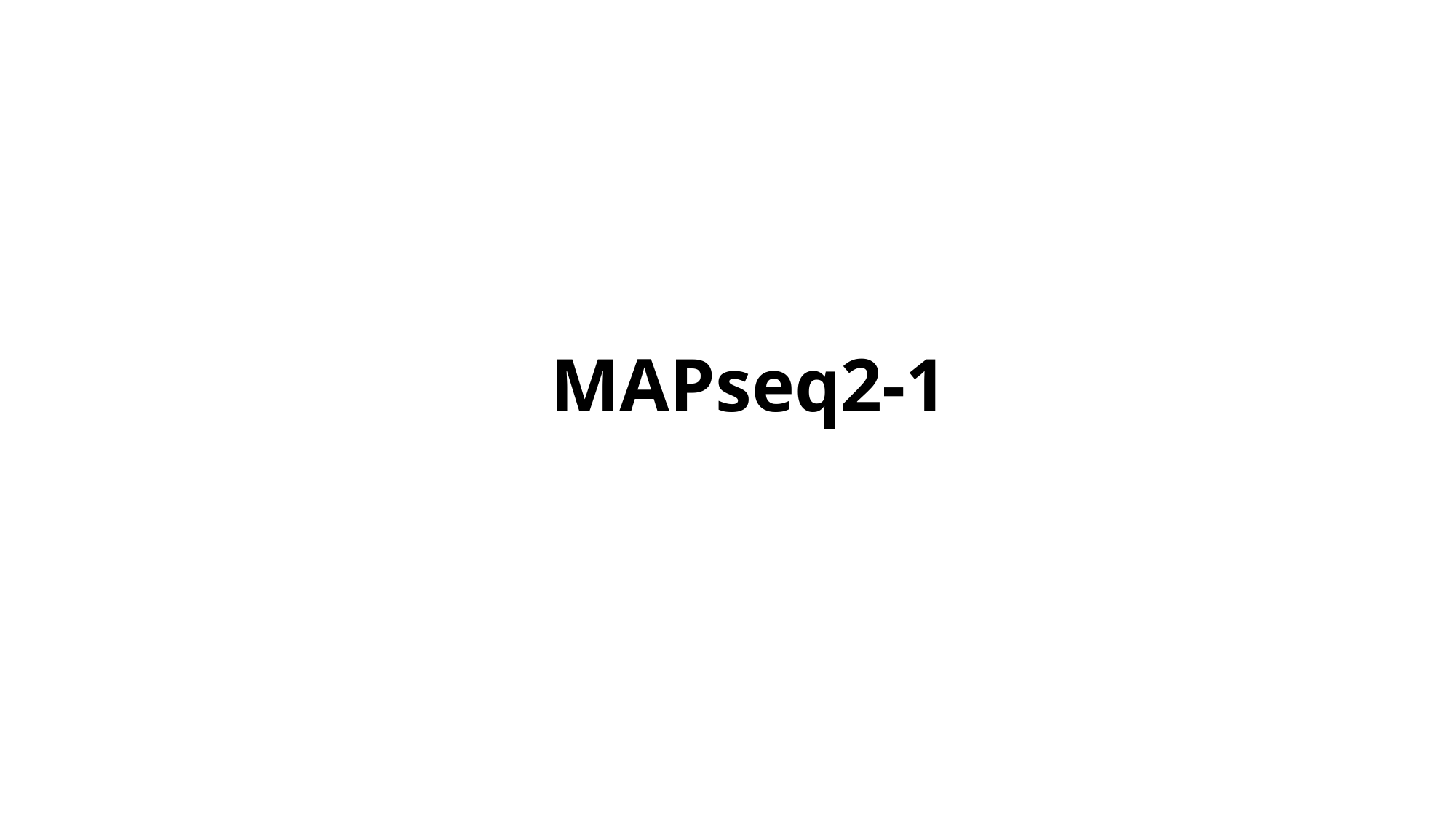

MAPseq2-1

### Slide 3
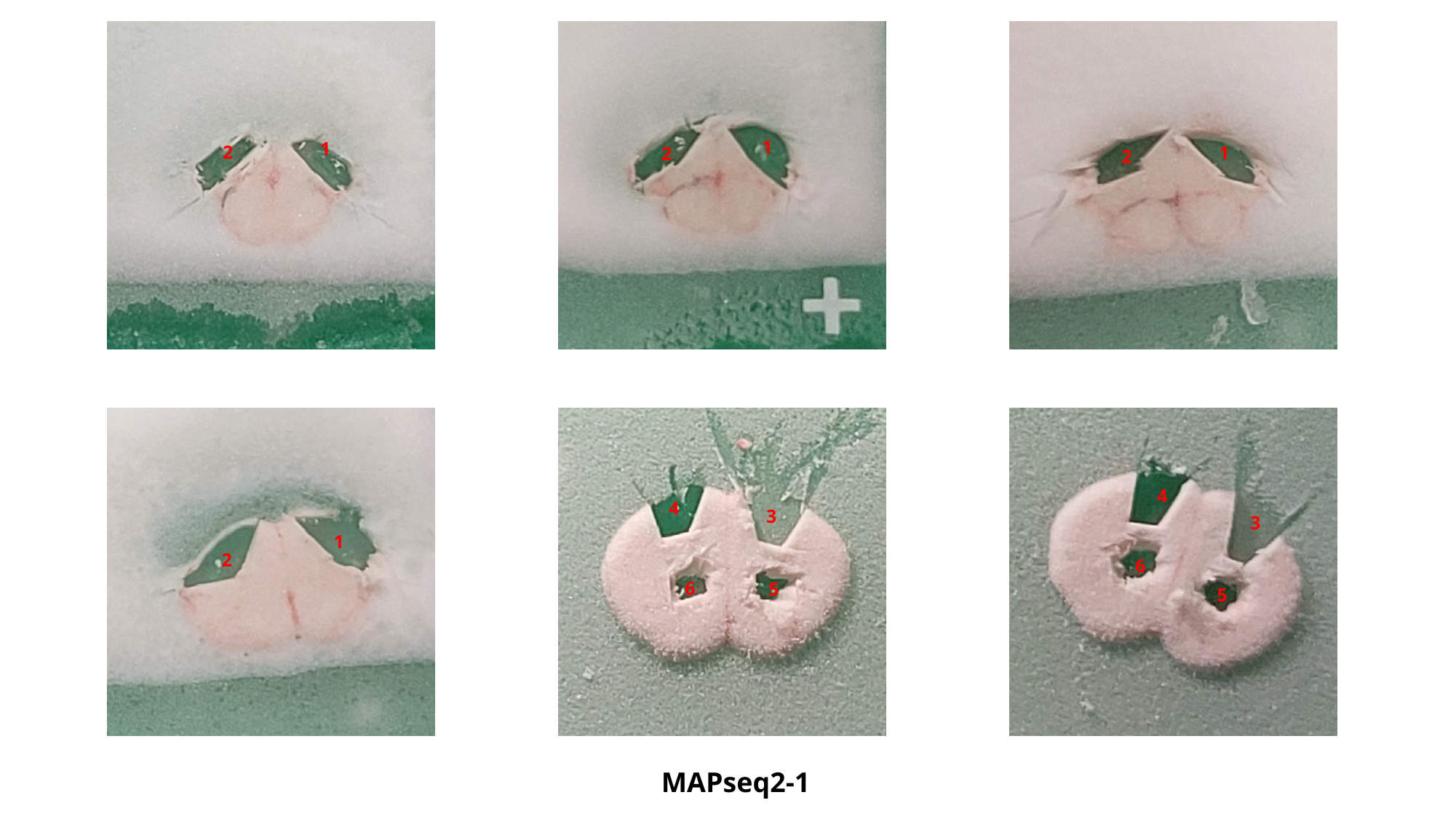

1
2
1
2
1
2
1
2
4
3
6
5
4
3
6
5
MAPseq2-1

### Slide 4
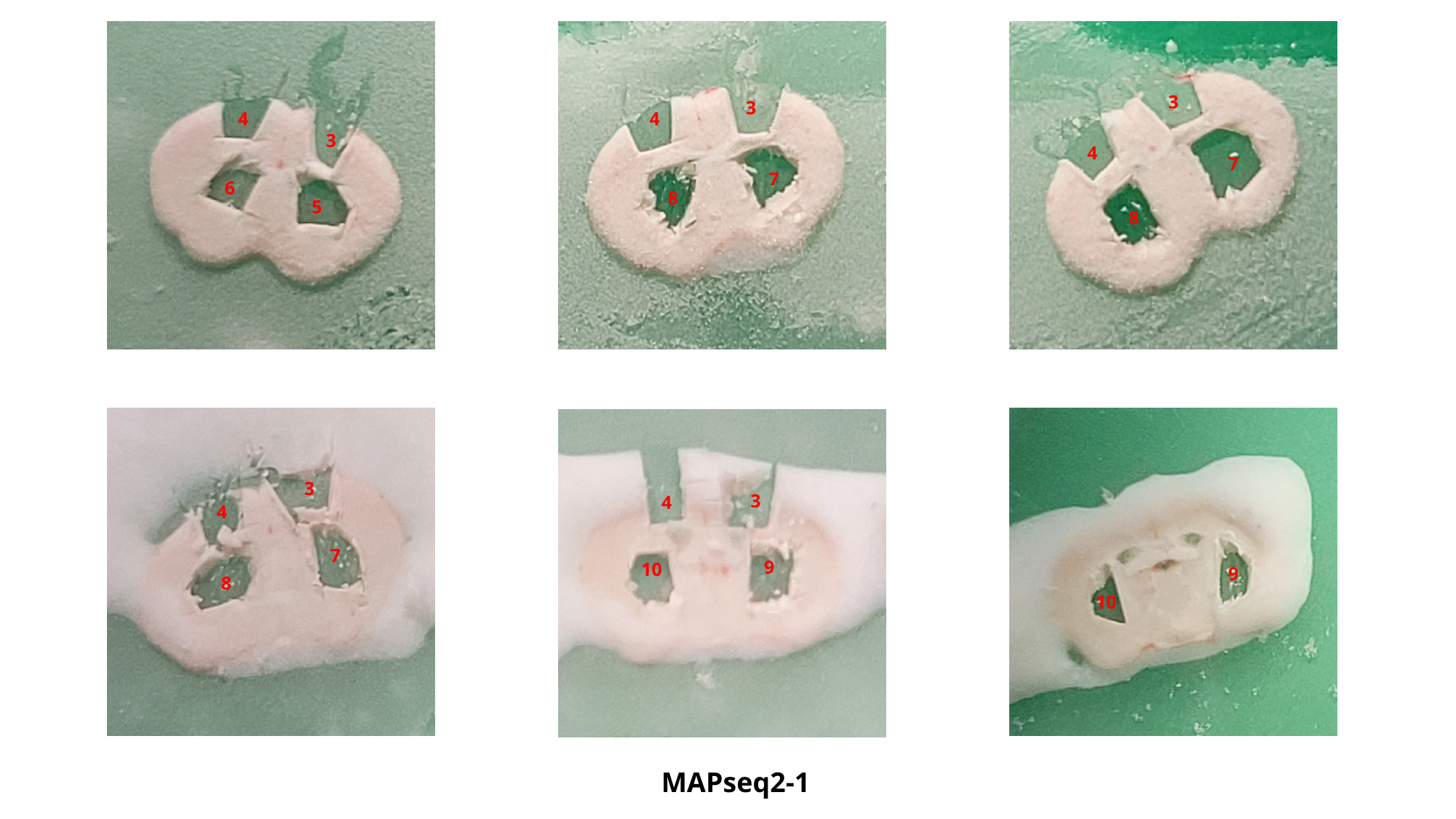

4
3
6
5
3
4
7
8
3
4
7
8
3
4
7
8
3
4
9
10
9
10
MAPseq2-1

### Slide 5
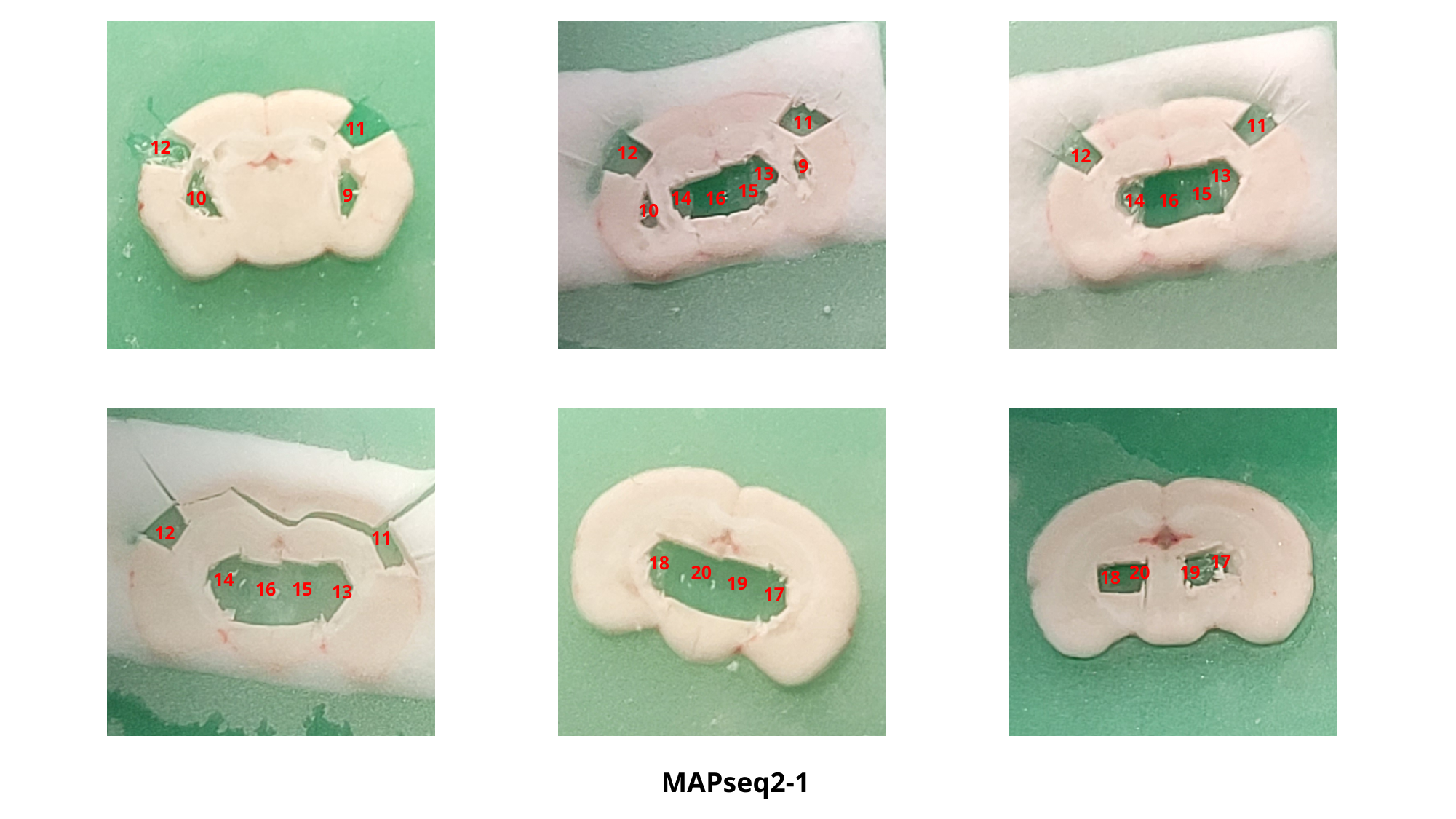

11
12
9
10
11
12
9
13
15
16
14
10
11
12
13
15
16
14
12
11
14
16
15
13
18
20
19
17
17
20
19
18
MAPseq2-1

### Slide 6
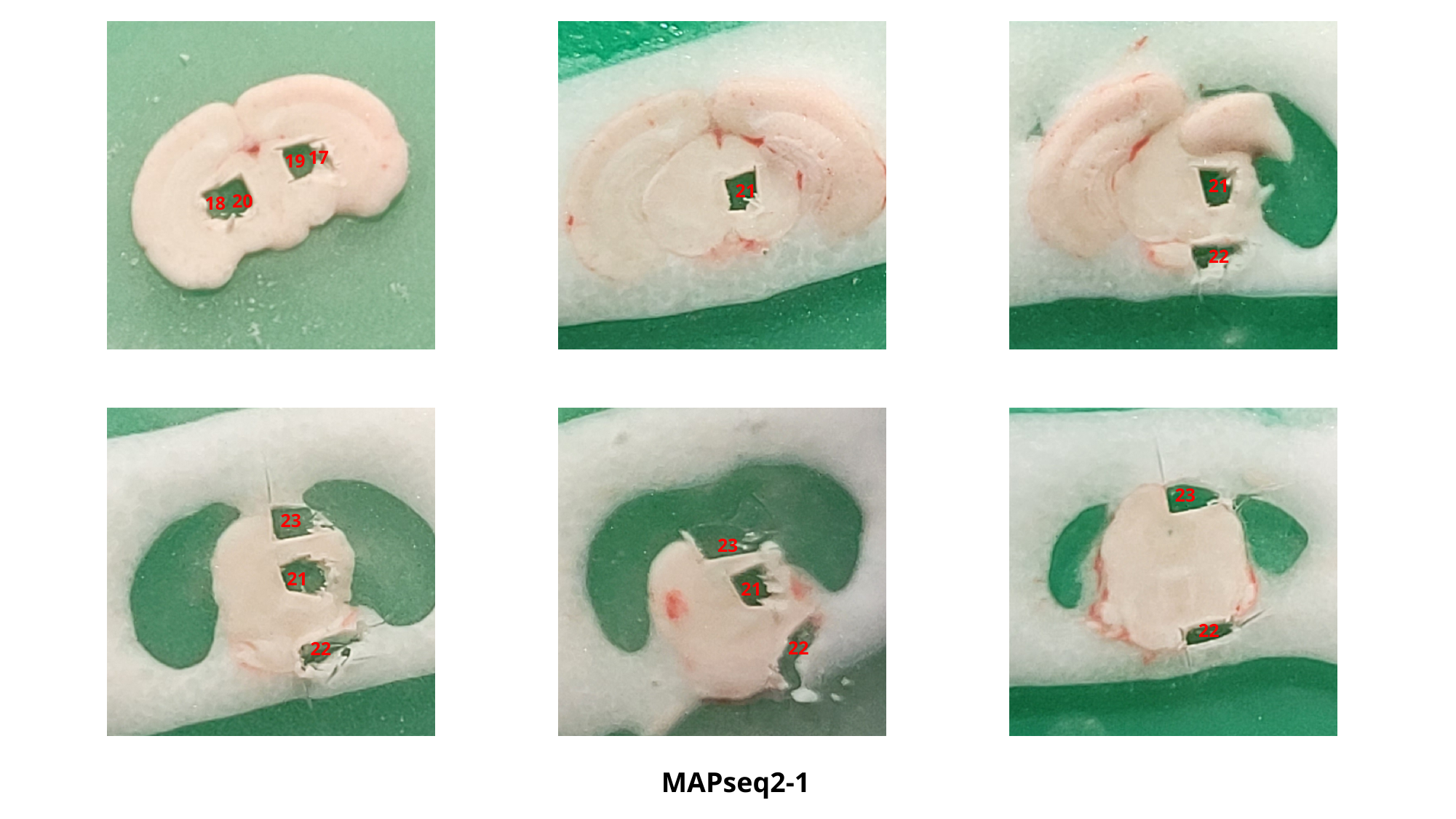

17
19
20
18
21
21
22
23
21
22
23
21
22
23
22
MAPseq2-1

### Slide 7
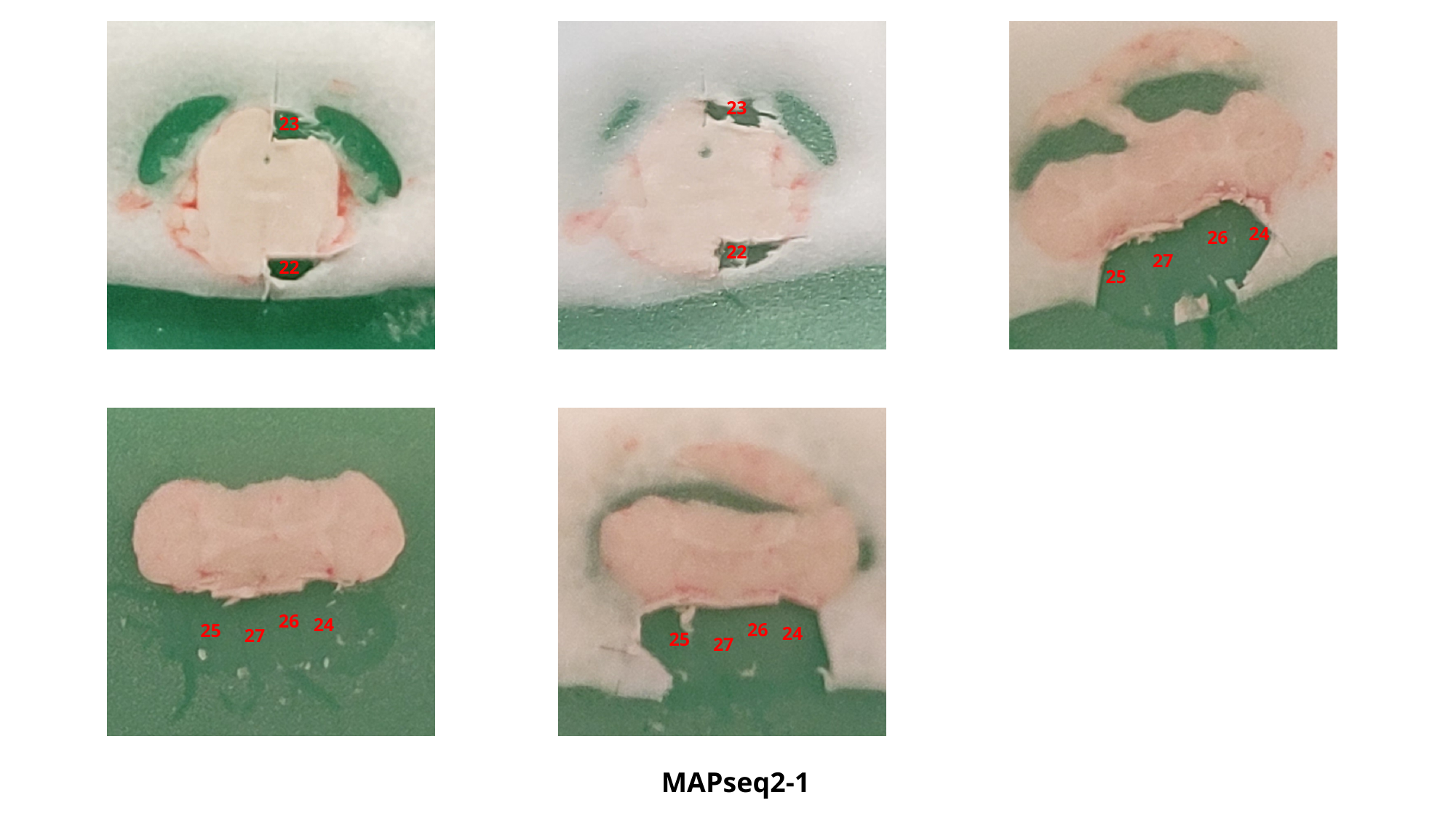

23
22
23
22
24
26
27
25
26
24
25
27
26
24
25
27
MAPseq2-1

### Slide 8
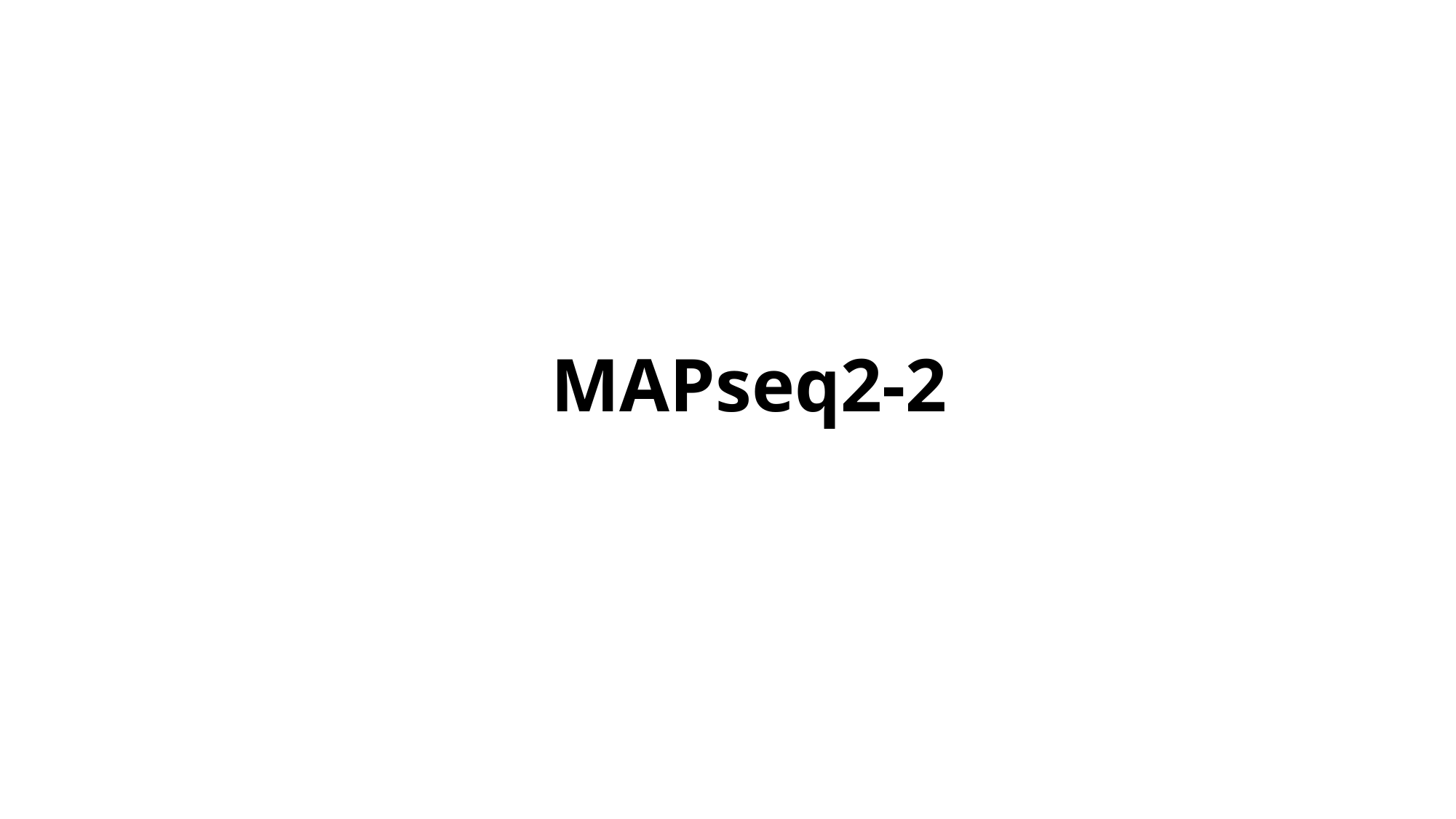

MAPseq2-2

### Slide 9
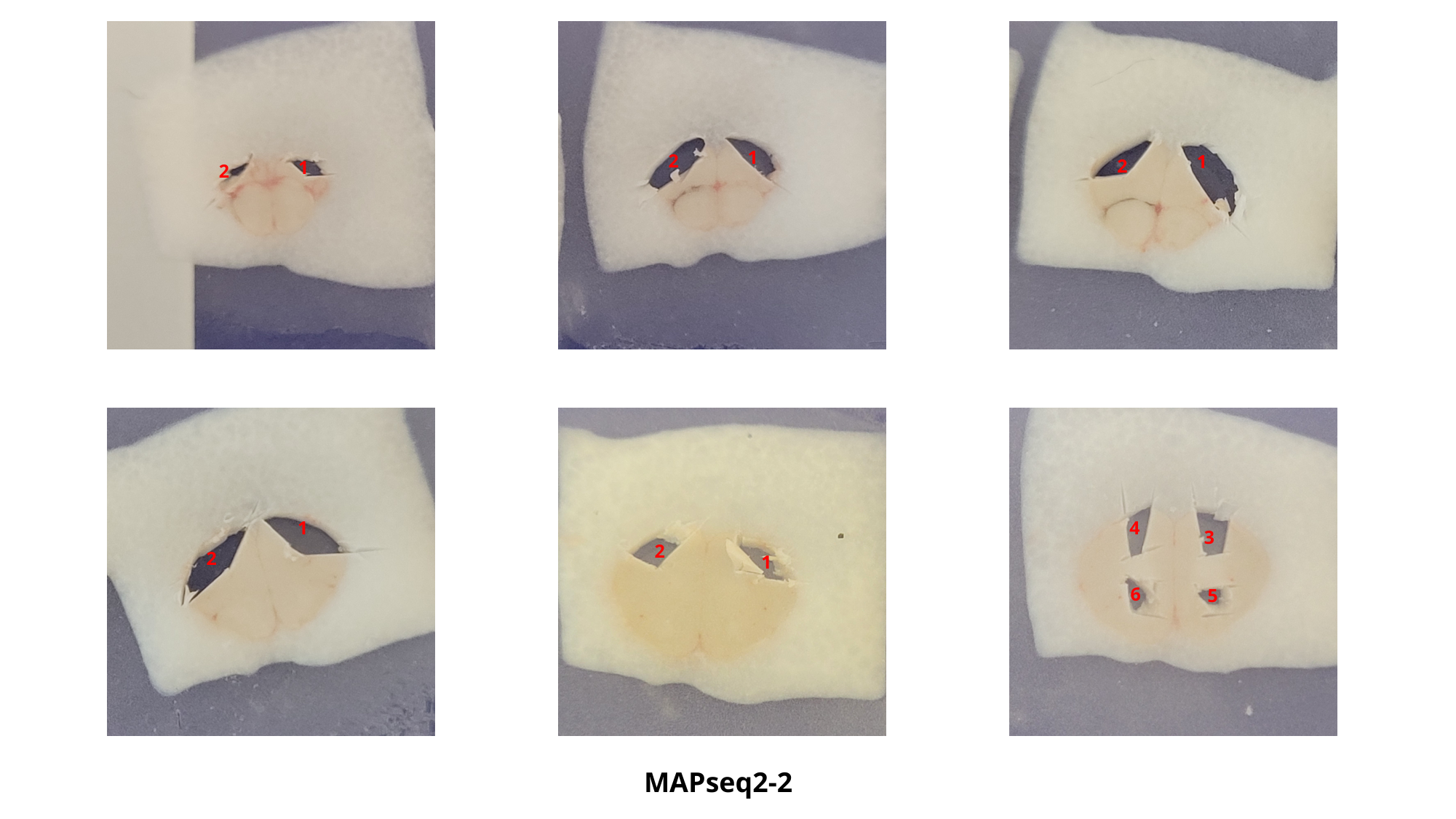

1
2
1
2
1
2
1
4
3
2
2
1
6
5
MAPseq2-2

### Slide 10
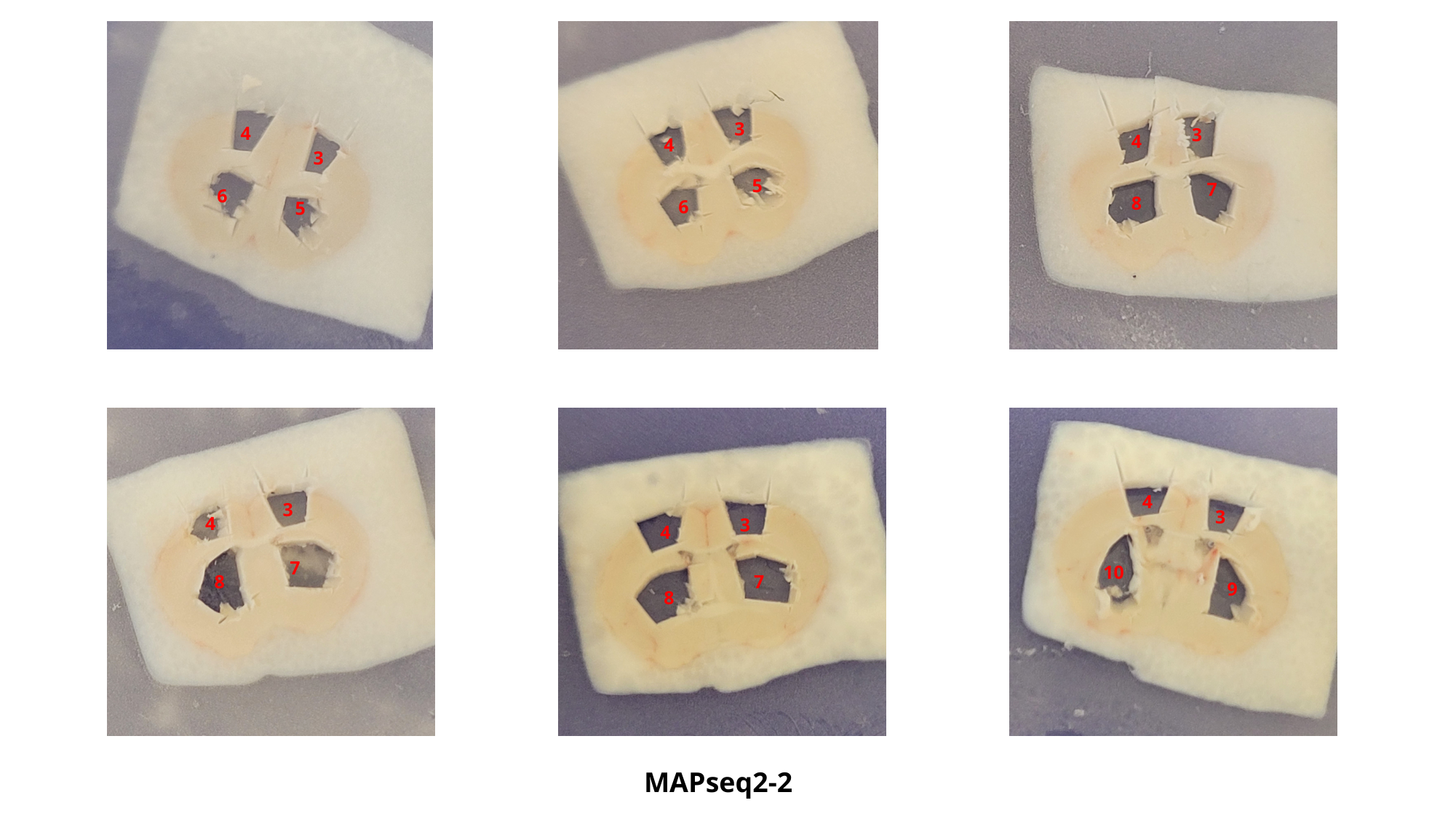

4
3
6
5
3
3
4
4
5
7
8
6
4
3
3
4
3
4
7
10
8
7
9
8
MAPseq2-2

### Slide 11
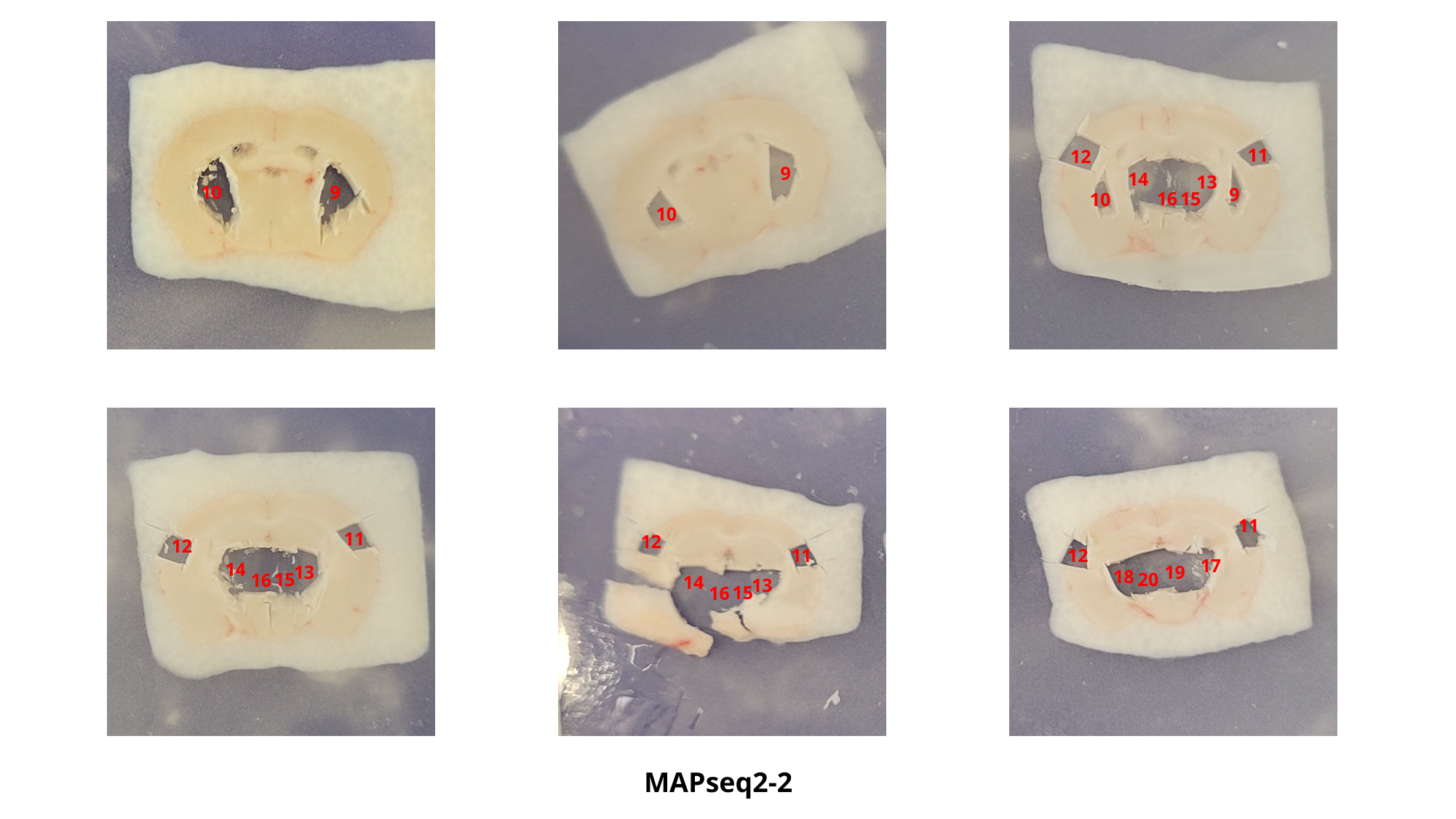

11
12
9
14
13
10
9
9
15
16
10
10
11
11
12
12
12
11
17
14
13
19
18
20
15
16
14
13
15
16
MAPseq2-2

### Slide 12
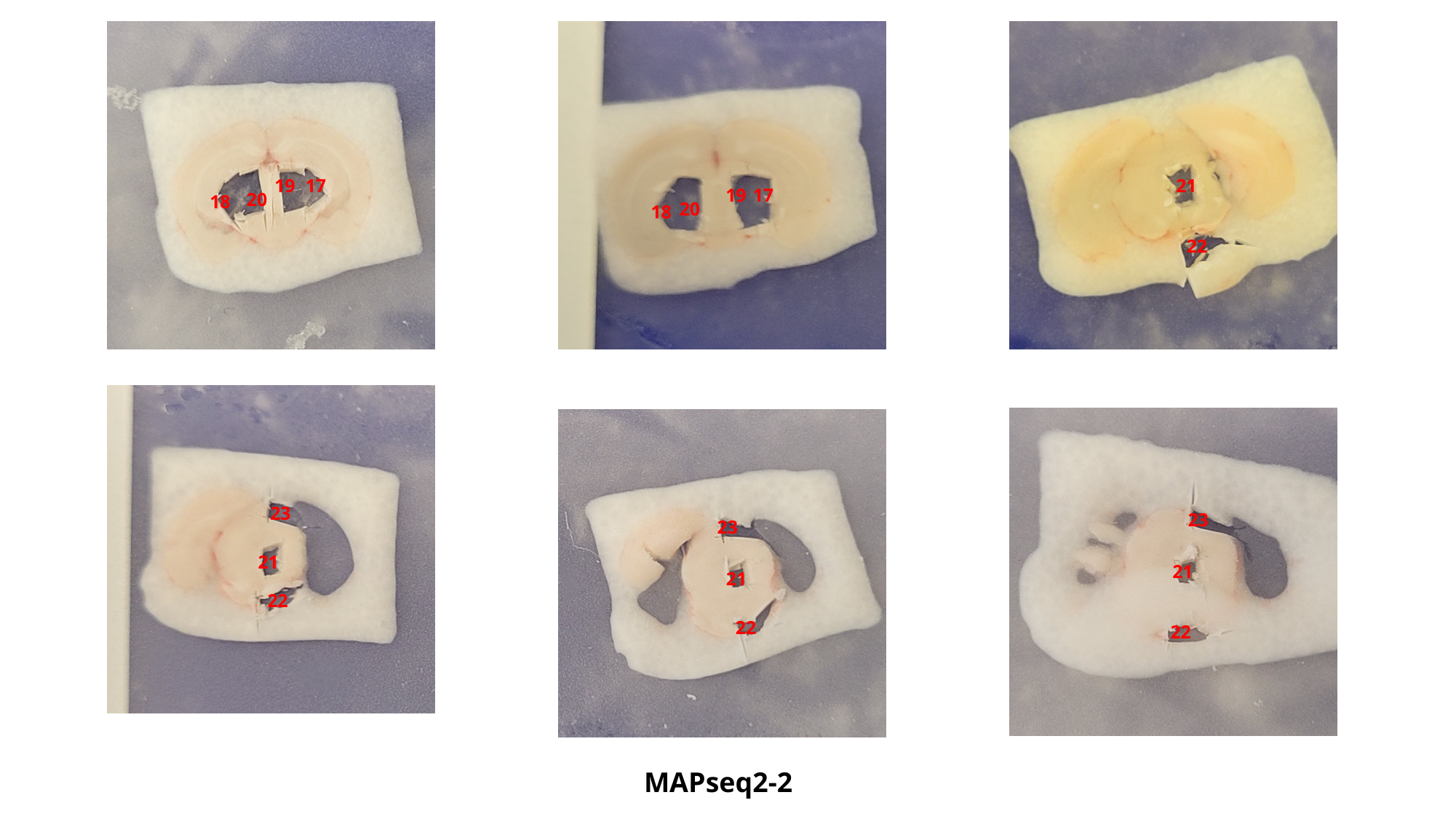

19
17
21
19
17
20
18
20
18
22
23
23
23
21
21
21
22
22
22
MAPseq2-2

### Slide 13
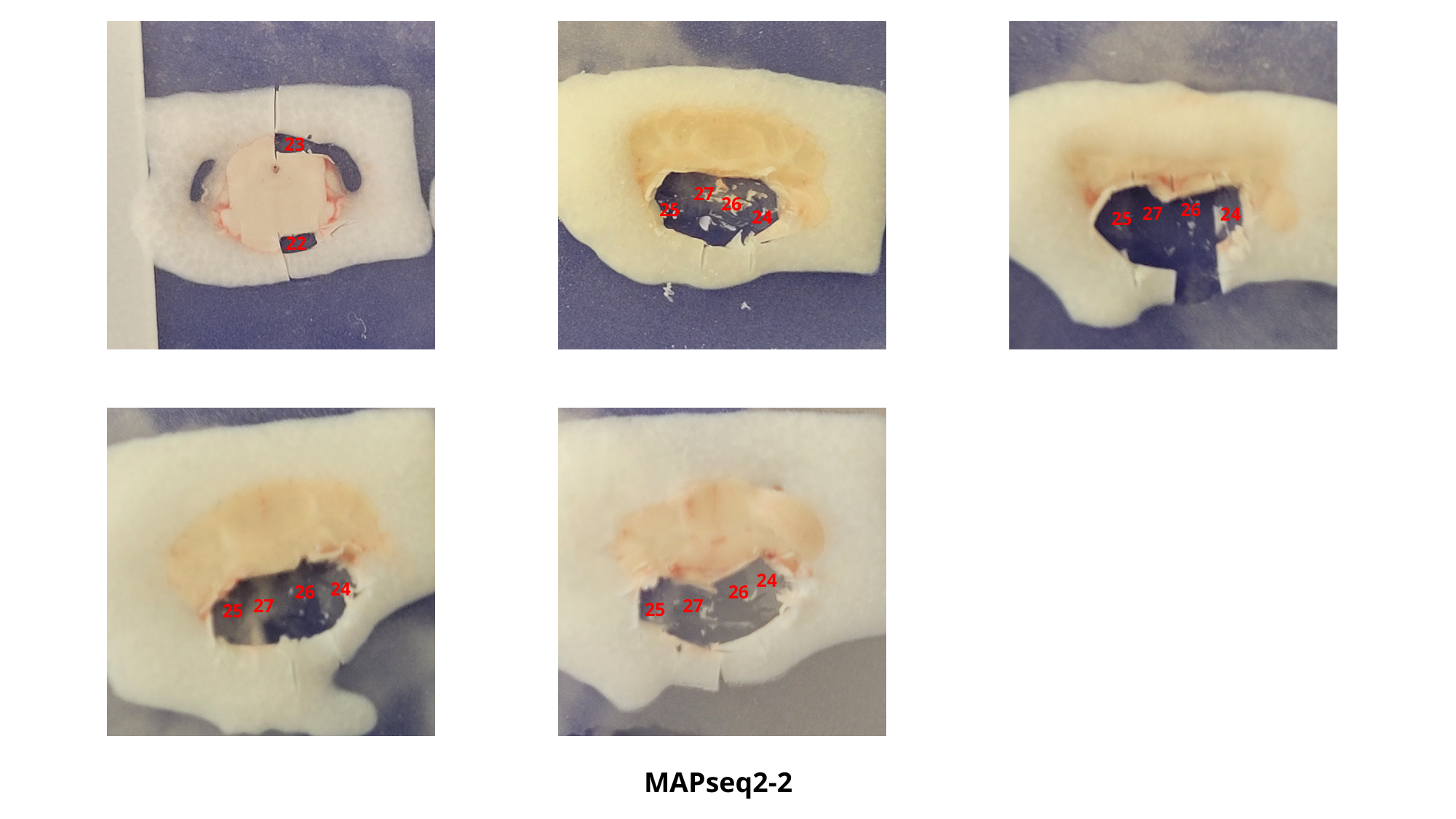

23
27
26
26
25
27
24
24
25
22
24
24
26
26
27
27
25
25
MAPseq2-2
